# Bacterial protein domains with a novel Ig-like fold target human CEACAM receptors

**DOI:** 10.1101/2020.06.09.141622

**Authors:** Nina M. van Sorge, Daniel A. Bonsor, Liwen Deng, Erik Lindahl, Verena Schmitt, Mykola Lyndin, Alexej Schmidt, Olof R. Nilsson, Jaime Brizuela, Elena Boero, Eric J. Sundberg, Jos A.G. van Strijp, Kelly S. Doran, Bernhard B. Singer, Gunnar Lindahl, Alex J. McCarthy

**Affiliations:** University Medical Center Utrecht, Utrecht University, Utrecht, The Netherlands; Institute of Human Virology, University of Maryland School of Medicine, University of Maryland, Baltimore, MD, USA; Department of Immunology & Microbiology, University of Colorado Anschutz Medical Campus, Aurora, CO, United States; Department of Biochemistry and Biophysics, Science for Life Laboratory, Stockholm University, Stockholm, Sweden; Institute of Anatomy, University Hospital, University Duisburg-Essen, Essen, Germany; Department of Pathology, Sumy State University, Sumy, Ukraine; Department of Medical Biosciences, Pathology, Umeå University, SE-901 85 Umeå, Sweden; Department of Laboratory Medicine, Division of Medical Microbiology, Lund University, Lund, Sweden; MRC Centre for Molecular Bacteriology & Infection, Imperial College London, London, United Kingdom; Department of Biochemistry, Emory University School of Medicine, Atlanta, GA, United States; Department of Chemistry, Division of Applied Microbiology, Lund University; SE-22100 Lund, Sweden

**Keywords:** Adhesin, immunoglobulin super-family, IgI, receptor, *Streptococcus agalactiae*

## Abstract

*Streptococcus agalactiae*, also known as group B *Streptococcus* (GBS), is the major cause of neonatal sepsis in humans. A critical step to infection is adhesion of bacteria at mucosal surfaces. Though several GBS adhesins have been identified, the host receptor targets of these adhesins remain unknown. We report here that surface-expressed β protein from GBS binds to human CEACAM1 and CEACAM5 receptors. A crystal structure of the complex showed that the IgSF domain in β represents a novel Ig-fold subtype called IgI3, in which unique features allow binding to CEACAM1. Bioinformatic assessments revealed that this newly identified IgI3 fold is not exclusively present in GBS. Instead, the IgI3 fold is predicted to be present in adhesins from other clinically important human pathogens. We confirmed the interaction between CEACAM1 and the predicted IgI3-containing adhesin in two different streptococcal pathogens. Overall, our results indicate that the IgI3 fold could provide a broadly applied mechanism for bacteria to target CEACAMs.

## Introduction

*Streptococcus agalactiae*, also known as Group B *Streptococcus* (GBS), is a major cause of pneumonia, septicaemia and meningitis in human neonates (Seale *et al*, 2017; Hall *et al*, 2017). Despite public health interventions, GBS is estimated to cause 410,000 infants infections and 150,000 stillbirths per year (Seale *et al*, 2017). Cellular adhesion to mucosal surfaces is the first critical step preceding infection (Patras & Nizet, 2018). Bacteria colonization is a multifactorial process that requires expression of adhesins that target extracellular matrix (ECM) constituents and/or host cell receptors (Pietrocola *et al*, 2018; Shabayek & Spellerberg, 2018). Indeed, GBS express a diverse array of surface adhesins and the exact repertoire varies from strain to strain, reflective of high plasticity of the GBS genome (Chen, 2019; Gori *et al*, 2020; Tettelin *et al*, 2005). Consequently, individual GBS strains differ in the host receptors that can be targeted during colonization. The mechanisms underpinning binding to ECM constituents are well-defined, and include targeting fibronectin, laminin and vitronectin (Deng *et al*, 2019; Spellerberg *et al*, 1999; Hull *et al*; Banerjee *et al*, 2011). In contrast, our knowledge of the mechanisms that GBS utilizes to directly promote host cell adhesion is limited (Patras & Nizet, 2018; Bolduc & Madoff, 2007; Pietrocola *et al*, 2018). In particular, the host receptors targets of several putative adhesins, including Rib and Sip proteins (Stålhammar-Carlemalm *et al*, 1993; Brodeur *et al*, 2000), remain uncharacterized. To develop vaccine or anti-bacterial strategies that interfere with mucosal colonization, insights into the full adhesin-receptor interaction repertoire is required (Pietrocola *et al*, 2018; Larsson *et al*, 1996; Heath, 2016; Michel *et al*, 1992).

We report that a subset of GBS strains interact with carcinoembryonic antigen-related cell adhesion molecules (CEACAM) receptors. Human CEACAMs are a family of 12 cell surface receptors belonging to the immunoglobulin (Ig) superfamily (IgSF) that are commonly expressed on epithelial cells, endothelial cells and leukocytes (Gray-Owen & Blumberg, 2006). CEACAMs regulate immune responses through formation of homophilic and heterophilic interactions (Bonsor *et al*, 2015b; Kuespert *et al*, 2006; Gray-Owen & Blumberg, 2006). These properties allow CEACAMs to regulate multiple physiological and pathophysiological processes including cell-to-cell communication, epithelial differentiation, apoptosis and regulation of pro-inflammatory reactions (Khairnar *et al*, 2018, 2015; Helfrich & Singer, 2019; Gray-Owen & Blumberg, 2006). Each CEACAM is composed of an N-terminal domain with V-set (IgV) fold, which confer the CEACAM-CEACAM interactions, and varying numbers of C2-set (IgC2) folds (Bonsor *et al*, 2015a, 2015b; Watt *et al*, 2001; Zhou *et al*, 1993). Interestingly, humans CEACAMs have been identified as docking receptors for Gram-negative bacteria (Javaheri *et al*, 2016; Brewer *et al*, 2019; Königer *et al*, 2016; Hill *et al*, 2001; Tchoupa *et al*, 2015; Chen & Gotschlich, 1996; Conners *et al*, 2008; Virji *et al*, 1996; Tchoupa *et al*, 2014). Therefore, we investigated the mechanism by which GBS binds to CEACAM receptors.

We report here that the GBS surface protein β binds specifically to human CEACAM1 and CEACAM5. Through structural methods and assays, we demonstrate that the IgSF domain in β protein contains unique structural features and represents a previously unrecognized variant of the IgI fold, which we termed the I3-set fold (IgI3). Homologs of β-IgI3 were identified in adhesins from several human bacteria including pathogens. We confirmed that CEACAM1 recognized one of these IgI3 variants and two streptococcal species, suggesting that the interaction between bacterial IgI3 domains and CEACAMs may be of general importance for host barrier colonization.

## Results

### The Group B *Streptococcus* surface β protein binds CEACAM1

GBS can successfully adhere to human mucosal cell lines, including vaginal, cervical and airway epithelial cell lines (Patras & Nizet, 2018). Yet, knowledge of the specific host factors that can be targeted by GBS for adhesion is lacking. As CEACAM receptors are mucosal docking targets for microorganisms (Tchoupa *et al*, 2014), we hypothesized that GBS interacts with CEACAM receptors. As individual GBS strains are likely to vary in the adhesion mechanisms due to genomic plasticity (Chen, 2019; Gori *et al*, 2020; Tettelin *et al*, 2005), we tested whether rCEACAM1 interacted with a panel of genetically diverse genome-sequenced GBS strains. We specifically observed binding to the GBS strains A909 and H36B (Fig. 1A, Supplementary Fig. 1A). To further test the specificity of the interaction, we also screened the ability of rCEACAM1 to interact with a variety of other Gram-positive species including *Streptococcus* spp., *Enterococcus* spp. and *Staphylococcus* spp. Notably, rCEACAM1 only bound to GBS strains included in our screen (Fig. 1A). This data reveals for the first time that GBS can bind to human CEACAM1 receptor.

**Figure 1:**
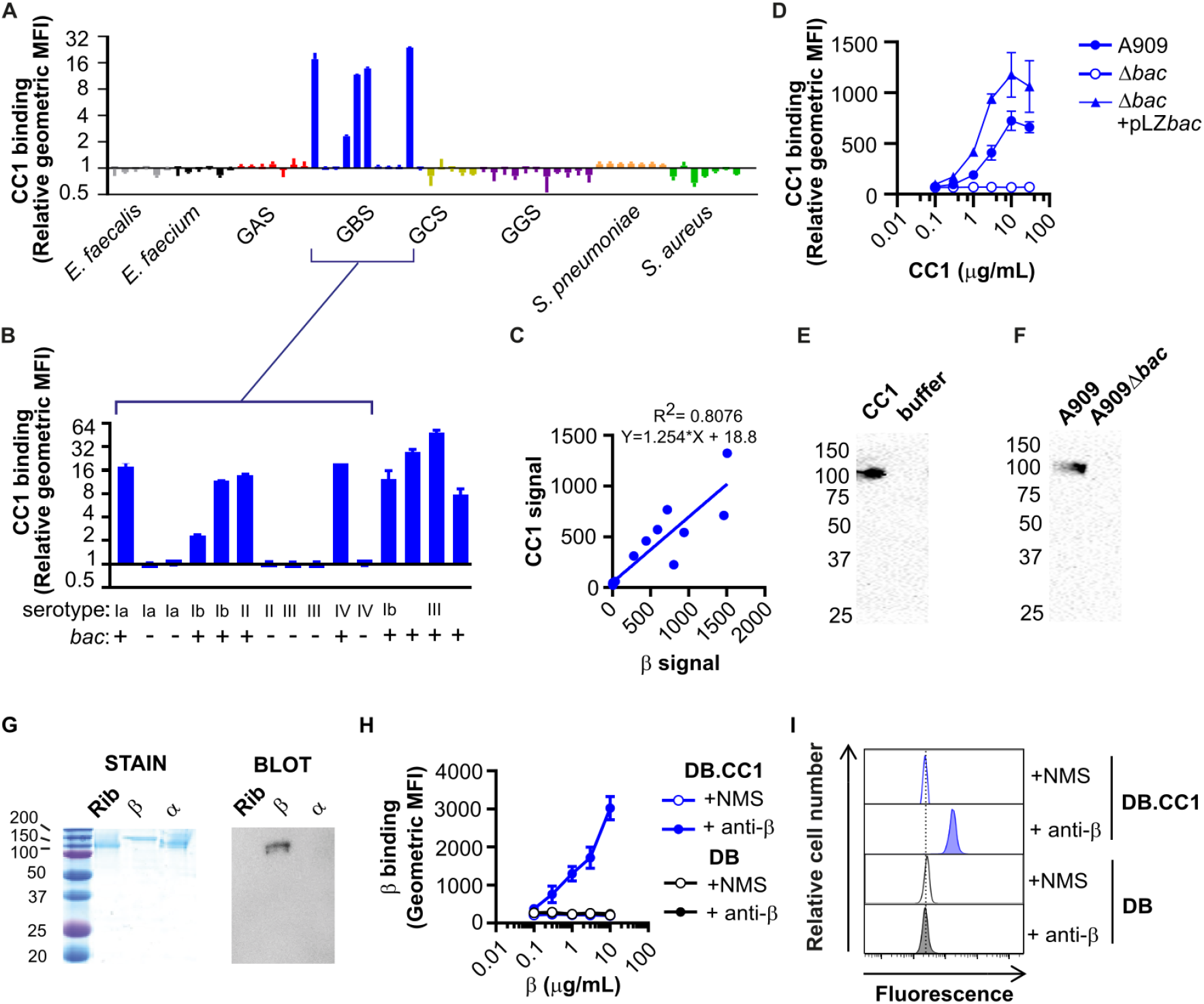
Human CEACAM1 binds to group B *Streptococcus* (GBS) via the surface-expressed β protein. **A and B)** Binding of recombinant (r)CEACAM1 (CC1)-HIS (10 μg/mL) to A) a panel of Gram-positive bacteria, namely *Enterococcus faecalis, Enterococcus faecium*, Group A *Streptococcus* (GAS), Group B *Streptococcus* (GBS), Group C *Streptococcus* (GCS), Group G *Streptococcus* (GGS), *Streptococcus pneumoniae*, *Staphylococcus aureus*, B) an expanded collection of GBS isolates, with serotype and carriage of *bac* gene shown. Mean and standard deviation (SD) values are reported for *n* = 3 independent experiments. **C)** Correlation between rCC1-binding capacity and β protein expression in GBS strains, respectively. β protein expression was quantified using rabbit anti-β protein serum and a secondary PE-conjugated goat anti-rabbit-IgG. rCC1 binding to live bacteria was quantified by a secondary anti-His-FITC monoclonal antibody (mAb). Mean binding values from *n* = 3 independent experiments. **D)** Concentration-dependent binding of rCC1-HIS to wild type (WT), Δ*bac* and Δ*bac* + pLZ.*bac* complemented A909 strains. Mean and SD values are reported for *n* = 3 independent experiments. **E)** Binding of human rCC1-HIS (30 μg/mL) to GBS strain A909 strain was analyzed by pull-down experiments followed by Western blot analysis. **F)** Human rCC1-HIS (30 μg/mL) binding to GBS A909 and A909Δ*bac* strains analyzed by pull-down experiments followed by Western blot analysis. **G)** Western blot analysis of rCC1-HIS (30 μg/mL) binding to purified β protein, but not to control GBS surface proteins α and Rib. Panel on the left show staining of proteins after separation by SDS-PAGE. **H** and **I)** Binding of purified β protein to dynabeads (DB) coated with rCC1 (DB.CC1) or control. Binding of β protein was detected using mouse anti-β serum or normal mouse serum (NMS), and PE-conjugated goat anti-mouse-IgG. H shows mean and SD values compiled from *n* = 3 independent replicates, and I shows representative flow cytometry plots using 10 μg/ml rCC1.

Next, we aimed to identify the GBS adhesin responsible for targeting rCEACAM1. We used a genome comparison study of Tettelin *et al*. (2005), to identify genes encoding cell-wall anchored proteins that were associated with rCEACAM1-binding phenotype. We identified that genomic island of diversity region 3.1 was present in rCEACAM1-binding strains (A909 and H36B) and absent in non-binding strains (515, COH1, NEM316 and NCTC10/84). This region encodes a cell wall-anchored protein known as β protein (encoded by *bac*). To test the hypothesis that β protein was responsible for CEACAM1 interaction, we screened rCEACAM1 binding to a broader collection of GBS isolates (Fig. 1B) and observed that CEACAM1 binding corelated with carriage of *bac* (Lindahl *et al*, 2005; Hedén *et al*, 1991). The intensity of rCEACAM1 binding differed between interacting GBS strains which may reflect differences in β expression. Indeed, β protein expression levels, detected using anti-β-serum (Supplementary Fig. 1B), correlated with rCEACAM1-binding capacity (Fig. 1C). Finally, we tested whether CEACAM1-binding was dependent on β protein expression. Deletion of *bac* in the GBS A909 reference strain abolished rCEACAM1 binding, whilst complementation with a vector carrying the *bac* gene re-established rCEACAM1 binding (Fig. 1D; Supplementary Fig. 1C). Mutation of other GBS surface adhesins or capsule, did not influence interaction with rCEACAM1 (Supplementary Fig. 1C). Additionally, we were able to pull down rCEACAM1 with GBS A909 (Fig. 1E), but not the Δ*bac* strain (Fig. 1F). Direct interaction of rCEACAM1 with purified β protein, but not with purified GBS surface proteins Rib or α, was demonstrated by Western blot analysis (Fig. 1G). Finally, β protein bound to CEACAM1-coated beads, but not control beads, in a concentration-dependent manner (Fig. 1H & 1I). Together, these data unequivocally show that GBS specifically binds to human CEACAM1 through expression of β protein.

### CEACAM1 binds to an IgSF domain in β protein

β protein is a multifunctional protein and interacts with several proteins of the human immune system, including factor H, Siglec-5, Siglec-7, Siglec-14 and IgA, through distinct domains (Fig. 2A) (Carlin *et al*, 2009; Fong *et al*, 2018; Ali *et al*, 2014; Hedén *et al*, 1991; Areschoug *et al*, 2002b; Lindahl *et al*, 1990; Nordström *et al*, 2011). To pinpoint the β protein domain interacting with CEACAM1, we expressed and purified recombinant β protein domains (B6N, IgABR, B6C, IgSF and β75KN) from *E. coli* (Fig. 2B) and tested their interaction with rCEACAM1. These domains include a part of β called IgSF because it adopts an immunoglobulin superfamily (IgSF) fold (Lindahl *et al*, 2005; Bateman *et al*, 1996). IgSF folds are composed of around 100 amino acids comprising two β sheets that pack face-to-face. CEACAM1 bound in a concentration-dependent manner to dynabeads (DB) coated with the biotinylated IgSF domain, but not to DB coated with other β protein domains or biotinylated-HSA (Fig 2C). As a control, we included rSiglec-5, which interacted only with DB coated with the B6N domain as described previously (Fig. 2C)(Nordström *et al*, 2011). To further test the interaction between CEACAM1 and the IgSF domain, we coated DB with rCEACAM1, which interacted with the IgSF domain, but not HSA, coupled to streptavidin (Fig. 2D). Thus, the IgSF domain in β specifically binds to human CEACAM1. Structural predictions of the β-IgSF domain suggest resemblance to a V-set Ig domain identified in SrpA, a cell-surface anchored protein in *Streptococcus sanguinis* (Fig. 2E)(Bensing *et al*, 2016). However, rCEACAM1 did not bind to the surface of a SrpA-expressing *S. sanguinis* strain (Supplementary Fig. 2). We denoted the IgSF fold in β as β-IgSF.

**Figure 2:**
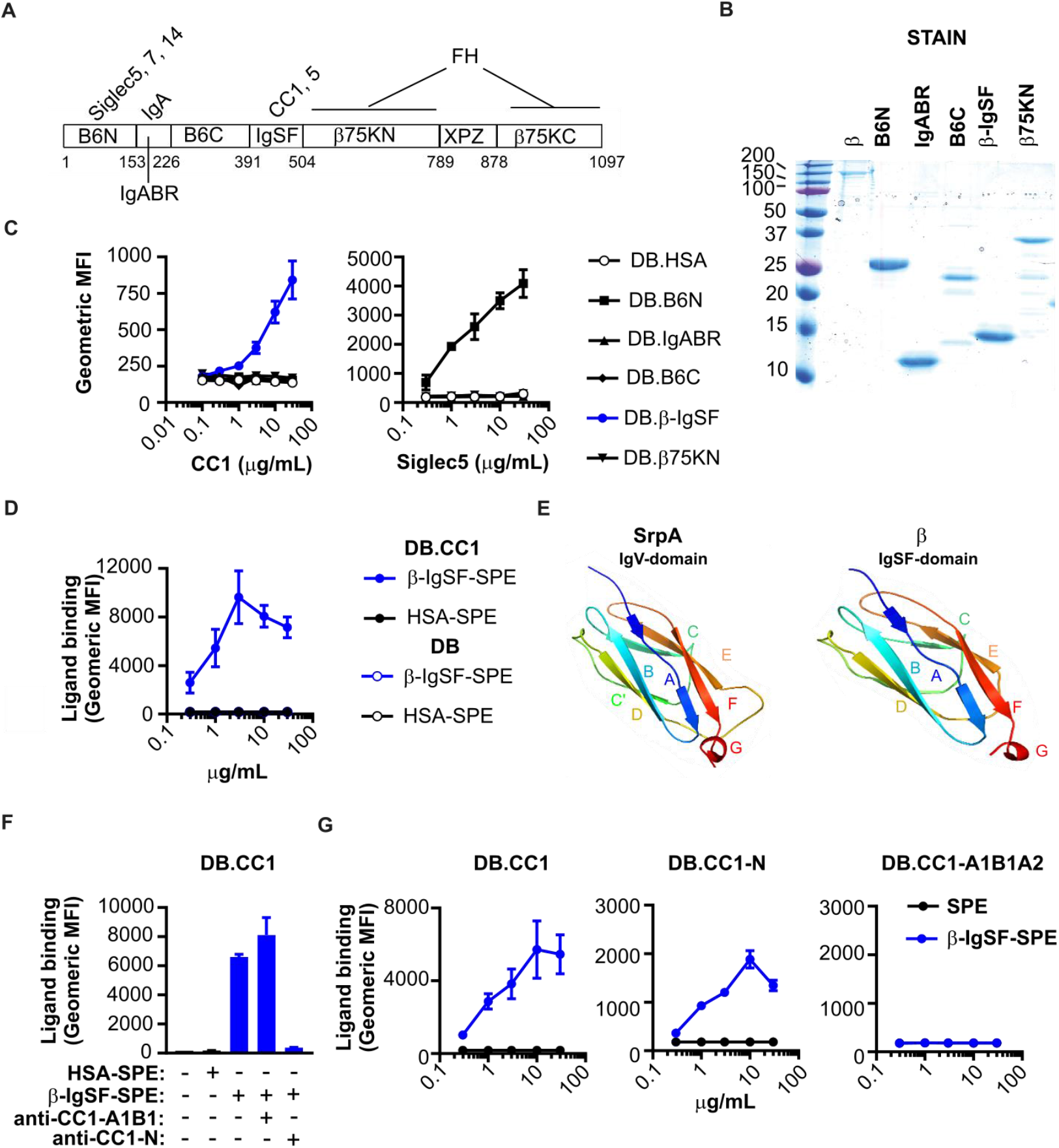
The IgSF domain in the β protein of GBS binds to the N-terminal domain of CEACAMs. **A)** Schematic representation of the β protein (Hedén *et al*, 1991). The first residue of each domain is indicated. XPZ is a proline-rich repeat region (Areschoug *et al*, 2002a) Known human ligand binding partners are indicated above each respective domain. IgA binds to a region denoted IgABR. Factor H binds the β75K region, but does not bind via XPZ. **B)** Staining of recombinant β protein domains (B6N, IgABR, B6C, IgSF and β75KN) after separation by SDS-PAGE. Recombinant proteins were expressed in *E. coli*. Purified β protein is used as a control. **C)** Binding of rCC1-His or rSiglec5-Fc to dynabeads (DB) coated with β protein domains. Mean and SD values are reported. **D)** Binding of varying concentrations of β-IgSF or human serum albumin (HSA) coupled to streptavidin-PE (SPE), to DB coated with CC1 (DB.CC1) or control protein (DB). Mean and SD values are reported. **E)** Structure of the relevant (Siglec-like sialic acid-binding) domain of the highest-scoring template 5KIQ (SrpA binding region), and predicted three-dimensional model of the IgSF domain from the β protein, constructed from the Hhpred alignment using MODELLER.(Webb & Sali, 2016) **F)** Inhibition of β-IgSF binding to DB.CC1 by anti-CC1-N mAb (clone CC1/3/5-Sab), but not by anti-CC1-A1B1 mAb (clone B3-17). β-IgSF-biotin was coupled to SPE. Mean and SD values are reported. **G)** Concentration-dependent binding of β-IgSF to DB coated with 100 μg/ml CC1 (DB.CC1), CC1-N (DB.CC1-N) or CC1-A1B1A2 (DB.CC1-A1B1A2). Assays utilized proteins coupled to SPE. Mean and SD values are reported.

CEACAM1 possesses four Ig-like domains (Supplementary Fig. 3A)(Gray-Owen & Blumberg, 2006), including the N-terminal IgV-like domain that forms the homo- and heterophilic interactions implicated in the CEACAM-mediated functions (Bonsor *et al*, 2015b). All bacterial ligands for CEACAM1 characterized to date target the N-terminal IgV-like domain. To identify the CEACAM1 domain targeted by β-IgSF, we tested whether domain-specific CEACAM1 monoclonal antibodies (mAb) could inhibit the interaction of β-IgSF with CEACAM1-coated DB. Only mAbs that block the N domain of CEACAM1 abolished the interaction with β-IgSF (Fig 2F; Supplementary Fig. 3B). Similarly, the *H. pylori* protein HopQ, which binds to the N-terminal IgV-like domain (Bonsor *et al*, 2018), blocked the β-IgSF-CEACAM1 interaction (Supplementary Fig. 3C). To test for direct interaction with the CEACAM1 N-terminal domain, we coated DB with rCEACAM1-N or rCEACAM1-A1B1A2 (Supplementary Fig. 3D) and observed concentration-dependent binding of β-IgSF only to DB coated with rCEACAM1-N (Fig. 2G). In the reciprocal experiment, rCEACAM1-N, but not rCEACAM1-A1B1A2, bound to DB coated with β-IgC2 (Supplementary Fig. 3E).

### The β-IgSF domains binds with high-affinity to human CEACAM1 and CEACAM5

Bacterial adhesins often bind to the N-terminal domain, not only of CEACAM1, but also CEACAM3, CEACAM5 and CEACAM6 reflecting the high sequence (~90% identity) and structural homology between CEACAM family members (Popp *et al*, 1999; Gray-Owen & Blumberg, 2006). However, no bacterial adhesins to date bind CEACAM8. To ascertain the CEACAM binding profile for β-IgSF, we measured the affinity of β-IgSF for unglycosylated N-terminal domains of these five CEACAMs, purified from *E. coli*, by isothermal calorimetry (ITC) (Fig. 3A; Supplementary Table 3). β-IgSF bound with high affinity to rCEACAM1-N (*K*_D_ = 96±2 nM) and rCEACAM5-N (*K*_D_ = 152±27 nM), but did not bind to rCEACAM3-N, rCEACAM6-N or rCEACAM8-N. The affinities of (β-IgSF)-(CEACAM1-N) and (β-IgSF)-(CEACAM5-N) interactions were comparable to those reported for other bacterial ligands (Bonsor *et al*, 2018; Korotkova *et al*, 2008). As the binding of β-IgSF is weaker to the isolated N-terminal domain compared to glycosylated CEACAM1 (Fig. 2G), other CEACAM1 domains may stabilise the interaction, as reported for the HopQ-CEACAM1 interaction (Bonsor *et al*, 2018). To confirm that the CEACAM binding profile of the protein is valid when expressed on the surface of GBS bacteria, we tested binding of the full-length glycosylated rCEACAMs to GBS by flow cytometry. The β-expressing GBS A909 strain recognized CEACAM1 and CEACAM5 only (Fig. 3B). Specificity for this subset was generally confirmed in a wider panel of β-expressing GBS strains (Supplementary Fig. 4), though variation existed in CEACAM5 binding. Although some β-expressing GBS strains did not bind CEACAM5, the β-IgSF domain displayed 100% sequence conservation amongst 57 GBS proteins identified through BLAST search (data not shown), indicating that intrastrain variation in CEACAM5 binding is most likely due to differential β expression.

**Figure 3:**
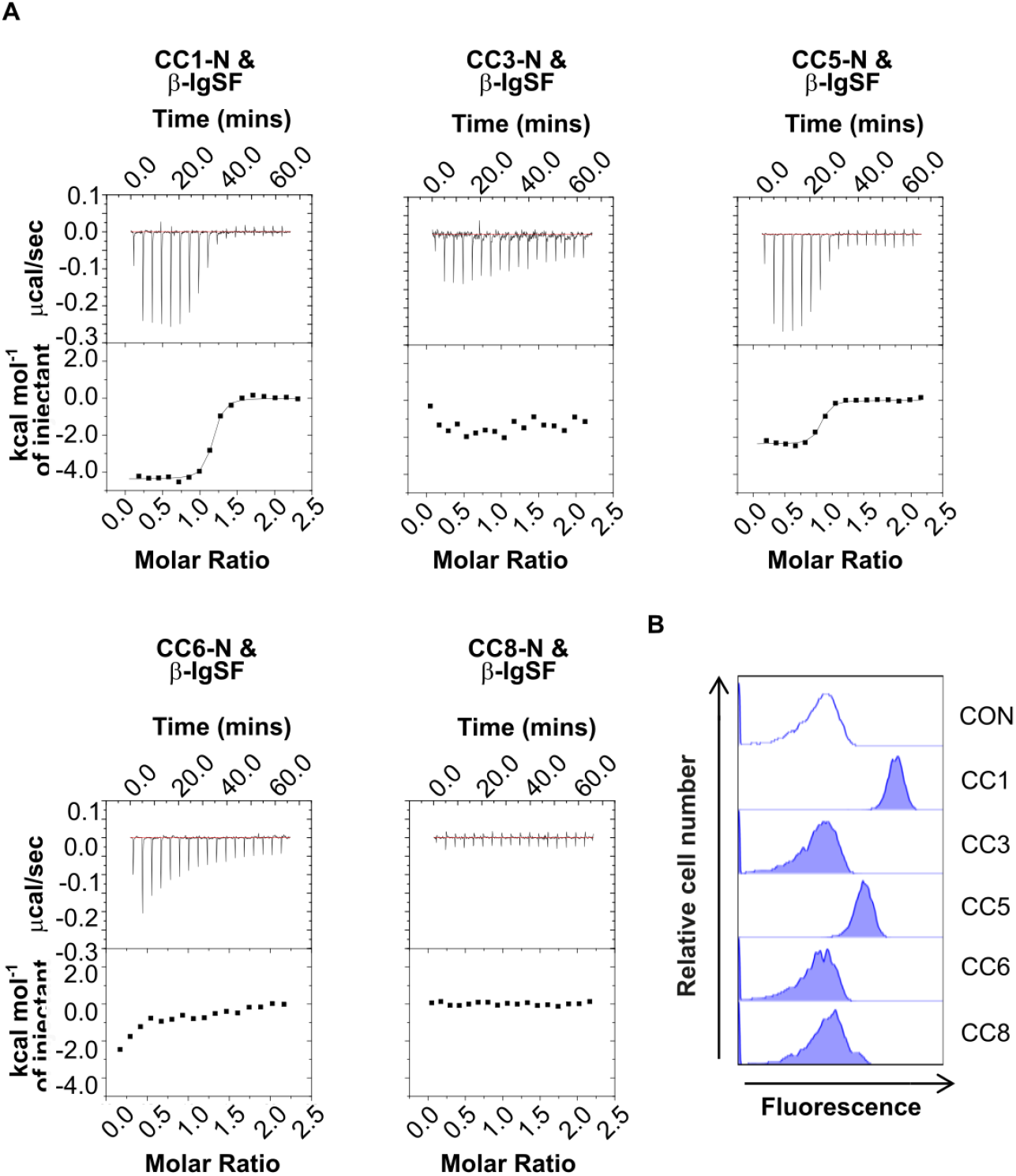
β-IgSF binds with high-affinity to human CEACAM1 and CEACAM5. **A)** Isothermal Titration Calorimetry (ITC) binding curves of CEACAM-N domains and β-IgSF. Experiments were performed using β-IgSF and N domains of (r)CEACAM1 (CC1), CEACAM3 (CC3), CEACAM5 (CC5), CEACAM6 (CC6) and CEACAM8 (CC8). **B)** Representative flow cytometry plots to analyze interaction of rCC1, CC3, CC5, CC6 or CC8 (10 μg/ml) with GBS A909 from *n* = 3 independent experiments.

### Expression of human CEACAMs on epithelial cells enhances GBS adhesion

As CEACAM engagement leads to enhanced cellular adhesion of other microorganisms, we hypothesized that CEACAMs could represent a novel cellular adhesion mechanism for GBS. To analyze whether β-expressing GBS can utilize human CEACAMs for cellular adhesion, we tested the binding of strain A909 to well-characterised CEACAM-expressing HeLa cells (Fig. 4A)(Bos *et al*, 1998). Consistent with rCEACAM binding, a higher percentage of the A909 inoculum was recovered from the CEACAM1- and CEACAM5-expressing HeLa cells in comparison to all other HeLa cell lines (Fig. 4A). Increased binding of GBS to human CEACAM1-expressing HeLa cells was confirmed by confocal microscopy (Fig. 4B). To ensure that the CEACAM-observed binding was not influenced by the background of the cell line, we assessed GBS adhesion to CEACAM-expressing CHO cells.(Hollandsworth *et al*, 2020) Also in this case, a higher percentage of the GBS inoculum was recovered from the CEACAM1- and CEACAM5-expressing CHO cells in comparison to all other CHO cell lines (Fig. 4C). For unknown reasons, there was considerable variation in GBS adhesion to the CEACAM1- and CEACAM5-expressing CHO cells. This variation could not be attributed to unstable CEACAM expression, as our cell lines expressed CEACAM at consistent levels (Supplementary Fig. 5A and 5B).

**Figure 4:**
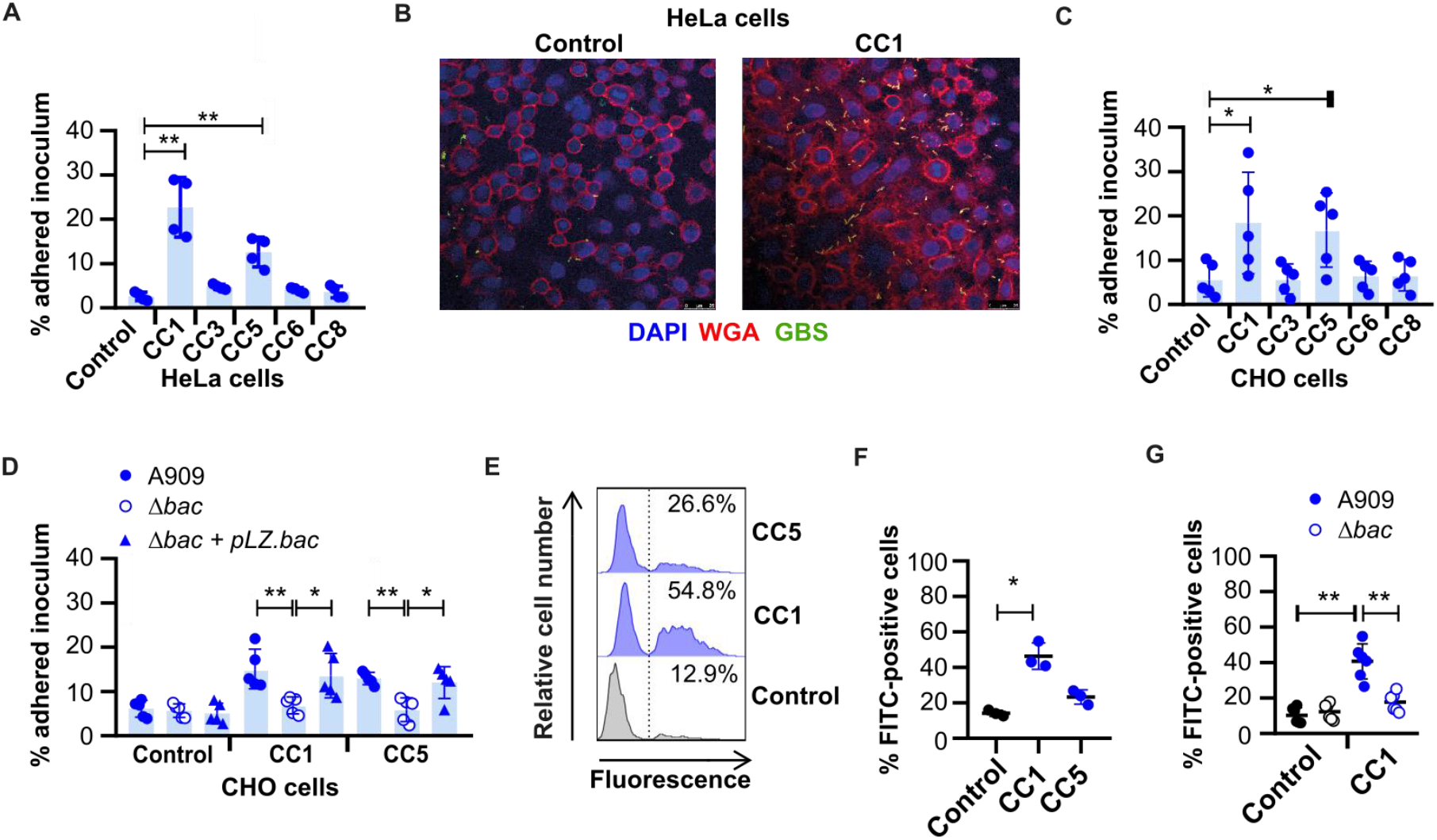
GBS exploit human CEACAM for enhanced adhesion to epithelial cells. **A)** Adherence of GBS A909 to human CEACAM1 (CC1)-, CEACAM3 (CC3)-, CEACAM5 (CC5)-, CEACAM6 (CC6)- or CEACAM8 (CC8)-expressing, or an empty vector control, HeLa transfectants at MOI of 10 for 30 mins. Mean and SD for *n* = 4 independent replicates are shown. **B)** Confocal imaging of FITC-labelled GBS A909 strain adhesion to human CC1-expressing or empty vector control HeLa transfectants. Cell membranes were stained with AF-647 conjugated wheat germ agglutinin (WGA)(red) and nuclei with DAPI (blue). **C)** Adherence of A909 to human CC1-, CC3-, CC5-, CC6- or CC8-expressing, or an empty vector control, CHO transfectants at MOI of 10 for 30 mins. Mean and SD from *n* = 5 independent replicates is displayed. **D)** Adhesion of WT, Δ*bac* and Δ*bac* + pLZ.*bac* complemented A909 strains to human CC1- or CC5-expressing CHO transfectants, or empty vector control cells. Mean and SD from *n* = 5 independent replicates is displayed. **E and F)** Adhesion of FITC-labelled A909 to human CC1-expressing, CC5-expressing or control CHO transfectants at an MOI of 10. Cell lines were detached and then incubated with FITC-labelled A909 for 30 mins at 4°C. Fluorescence of cells was measured by flow cytometry analysis. The percentage of fluorescent cells was calculated for each population. Representative flow cytometry plots are shown E, and the integrated results displaying the mean and SD from 3 independent replicates are reported in F. **G)** Adhesion of FITC-labelled A909 strains to human CC1-expressing or control CHO transfectants at MOI of 10. Cell lines were detached and then incubated with FITC-labelled A909 strains for 30 mins at 4°C. Fluorescence of cells was measured by flow cytometry analysis, where the percentage of fluorescent cells was calculated. Mean and SD for each population for *n* = 6 independent replicates is displayed. Data in A, C, D, F and G was analyzed by repeated One-way ANOVA with Sidak’s multiple comparisons. \**P*<0.05, \*\**P*<0.01.

To confirm that adhesion of the GBS strain A909 to the CEACAM1-expressing CHO cell was dependent on β protein, we also assessed binding of the *bac* deletion mutant and complemented strain in this same system. GBS adherence was abolished by mutation of the *bac* gene, and the phenotype could be recapitulated by a complemented mutant (Fig. 4D). This result was confirmed by confocal microscopy (Supplementary Fig. 5C). In agreement with the results obtained with purified proteins (Fig. 2F), the binding of β-expressing GBS to CEACAM1-expressing CHO cells was inhibited by blocking the CEACAM1-N domain with a specific mAb (Supplementary Fig. 5D). Moreover, pre-incubation of A909 with rCEACAM1-N, but not rCEACAM8-N, impaired adhesion to CEACAM1-expressing CHO transfectants (Supplementary Fig. 5E). This indicates that the CEACAM1-N and β-IgSF domains were responsible for the cellular adhesion phenotype.

To rule out the possibility that differences in adhesion of A909 wildtype and Δ*bac* strains to CHO cells reflected interstrain variation in growth rates during adhesion at 37°C, we developed an alternative assay in which we assessed adhesion of FITC-labelled GBS strains to detached cell lines during incubation at 4°C. A higher percentage of CEACAM1-expressing CHO cells were FITC-positive upon incubation with A909 in comparison to control cells (Fig. 4E and 4F). However, this was not observed for CEACAM5-expressing CHO cells, likely because the N domain of CEACAM5 was cleaved during the detachment process as demonstrated by the inability of an anti-CEACAM5-N mAb to bind to these cells (Supplementary Fig. 5A and 5B). As expected, adhesion of FITC-labelled A909 to the CEACAM1-expressing CHO cell was abolished by mutation of *bac* (Fig. 4G; Supplementary Fig. 5F). Collectively, the data therefore indicate that binding to human CEACAM receptors, via β, is a novel cellular adhesion mechanism for GBS.

### Crystallography reveals that the IgSF domain in β adopts a novel Ig fold, the IgI3 fold

To gain insights into β-IgSF binding mechanisms, we aimed to solve its structure. We solved the structure in complex with CEACAM1-N at a resolution of 3.0Å (Fig. 5A). The asymmetric unit contains two molecules of the (β-IgSF)-(CEACAM1-N) complex. The β-IgSF domain has the characteristic features of an Ig fold, principally a pair of β sheets built of anti-parallel β strands that surround a hydrophobic core (Fig. 5B). IgSF domains can be classified into (variable) V-set, (constant) C-set or (intermediate) I-set, with differentiation based on the number and placement of β-strands between the conserved cysteine residue disulphide bridge (Fig. 5C)(Wang & Springer, 1998; Wang, 2013). The V-set Ig domain contains ten β strands with four strands found on one sheet (*ABED*) and six strands on the other (*A’ GFCC’C”*). C-set Ig domains lack the *A’* and *C”* strands, and are further grouped into C1 or C2 based on presence or absence of the *D* strand, respectively. I-set Ig domains lack the *C”* strand and are classified into I1 or I2 based on presence or absence of a *D* strand, respectively. The β-IgSF domain has two β-sheets labelled *ABED* and *AGFC*, with sheets connected by the *BC*, *EF*, *CD* and *AA’* loops (Fig. 5B). Therefore, β-IgSF has an I-set fold topology that most closely resembles an I1-set domain (Fig. 5C). However, β-IgSF lacks cysteines and disulfide bridges that are characteristic for I-set folds. Furthermore, the β-IgSF domain possesses a truncated *C* strand that is directly followed by 1.5-turn α-helix (Fig. 5C). These features were not observed in the structurally characterised IgI1 domains in the DALI or PDBeFOLD databases that β-IgSF most closely resembled, including macrophage colony stimulating factor 1 (MCS-F) and intracellular adhesion molecules 3 (ICAM-3) (Fig. 5D, 5E & 5F). Therefore, the topology of β-IgSF domain represents a previously unrecognized IgI fold subtype, denoted here as I3-set Ig (IgI3) domain. Accordingly, the domain in β will now be referred to as β-IgI3. The unique features of this domain are i) the absence of cysteine residues, ii) absence of *C’C”*, and iii) a truncated *C* that is directly followed by a 1.5-turn α-helix. Of note, the unique β-IgI3 region stretching from *C* to *D* strand possesses protruding hydrophobic residues, such as F42, located between *C* and α-helix, L46 located in the α-helix, and V53 located in the *CD* loop (Fig 5G).

**Figure 5:**
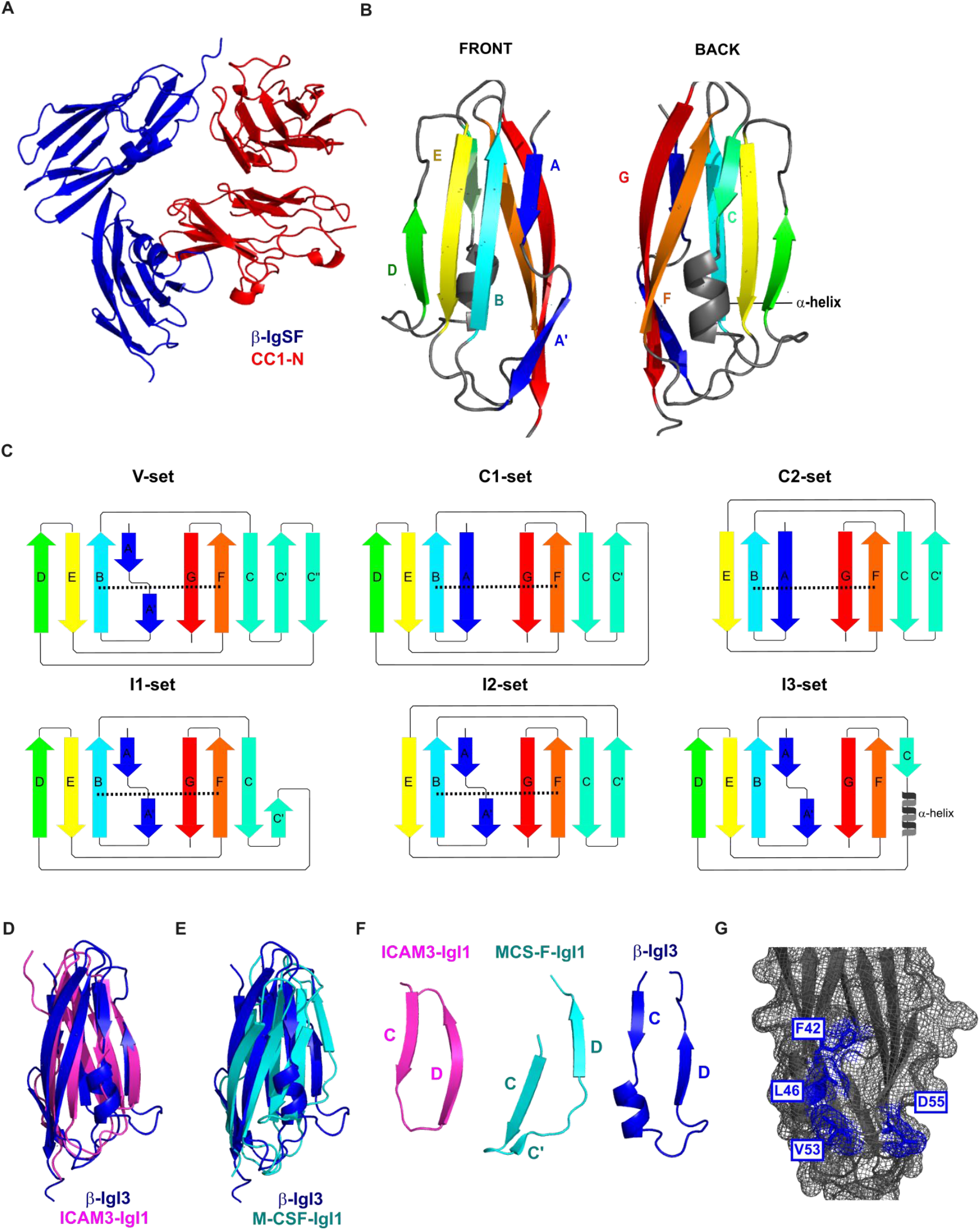
The Ig-like domain of β protein represents a new IgI fold subtype. A) Dimer co-crystal structure of β-IgSF (blue) and CEACAM1-N (CC1-N; red). **B)** Molecular structure of β-IgSF, in which each β-strand is coloured differently. The 1.5-turn α-helix shown in grey is located between the *C* strand and the *D* strand. **C)** Illustration of the secondary structure organisation of the V-set, C1-set, C2-set, I1-set, I2-set and I3-set domains. Each secondary structure is composed of varying combinations of the β-strands *A, A’, B, C, C’, C”, D, E, F, G*. Disulfide bridges betwen the *B* strand and *F* strand are represented as a black dashed lines. The C-set domains lack the *A’* and *C*” strands, and are subtyped into C1-set and C2-set based on presence and absence of the *D* strand, respectively. The I-set domains lack *C*” strand, and are subtyped into C1-set and C2-set based on presence and absence of the *D* strand, respectively. The IgSF fold from β protein has the I3-set domain, characterised by absence of disulfide bridge, presence of the *D* strand, and presence of a truncated *C* strand followed by a 1.5-turn α-helix. **D** and **E)** Superposition of the IgI3 domain from β protein (blue) onto IgI1 domains from ICAM3 (magenta, PDB entry 1T0P; D) or macrophage colony stimulating factor 1 (MCS-F; turquoise, PDB entry 3EJJ; E). **F)** Comparison of β-IgI3 (blue) with the most similar folds; the IgI1 domain of MCS-F (turquoise) and the IgI1 domain 1 of ICAM-3 (magenta,). **G)** The molecular surface mesh of β-IgI3 residues showing protruding hydrophobic residues located between the *C* and *D* strands in blue.

### Identification of interacting residues in the (β-IgI3)–(CEACAM1-N) complex

CEACAMs form their homophilic and heterophilic interactions through interactions promoted by the *A’GFCC’* face in the N-terminal V-set domains (Fig. 5A and Fig. 6A). Bacterial ligands also bind to the *A’GFCC’* face (Brewer *et al*, 2019; Conners *et al*, 2008; Villullas *et al*, 2007; Korotkova *et al*, 2008; Bonsor *et al*, 2018; Virji *et al*, 1999; Moonens *et al*, 2018). The *H. pylori* ligand HopQ uses a coupled folding and binding mechanism (Bonsor *et al*, 2018), and simulated docking indicated that the *E. coli* ligand AfaE binds to the CEACAM dimerization interface through the *BE* strands and *DE* loop of an incomplete Ig fold (Korotkova *et al*, 2008; Anderson *et al*, 2004). The structure determination by X-ray crystallography (Fig. 6B; Supplementary Table 4) of the complex of β-IgI3 and CEACAM1-N at 3 Å resolution showed that β-IgI3 binds to the *A’GFCC’* face of CEACAM1 through residues located in the *C* to *D* strand region including the α-helix (Fig. 6C). Fig. 6B shows the interacting residues on the surfaces of the two molecules. Specifically, L46 in the α-helix of β-IgI3 interacts by van der Waals forces with F29 and L95 (distances 3.11Å and 3.61Å) located in the *C* and *G* strands of CEACAM1, respectively (Supplementary Fig. 6A; Supplementary Table 5). In addition, the protruding β-IgI3 residue V53 is within close contact of I91 (distance 3.16Å, *F* strand) and S32 (distance 3.36Å, *C* strand) of CEACAM1, D55 is within close contact of Y34 (*C* to *C’* loop) of CEACAM1, and F42 is in close contact of L95 (distance 2.87Å, *G* strand) of CEACAM1 (Supplementary Table 5). No conformational changes are observed in CEACAM1 upon binding of β-IgI3 (Supplementary Fig. 6B).

**Figure 6.**
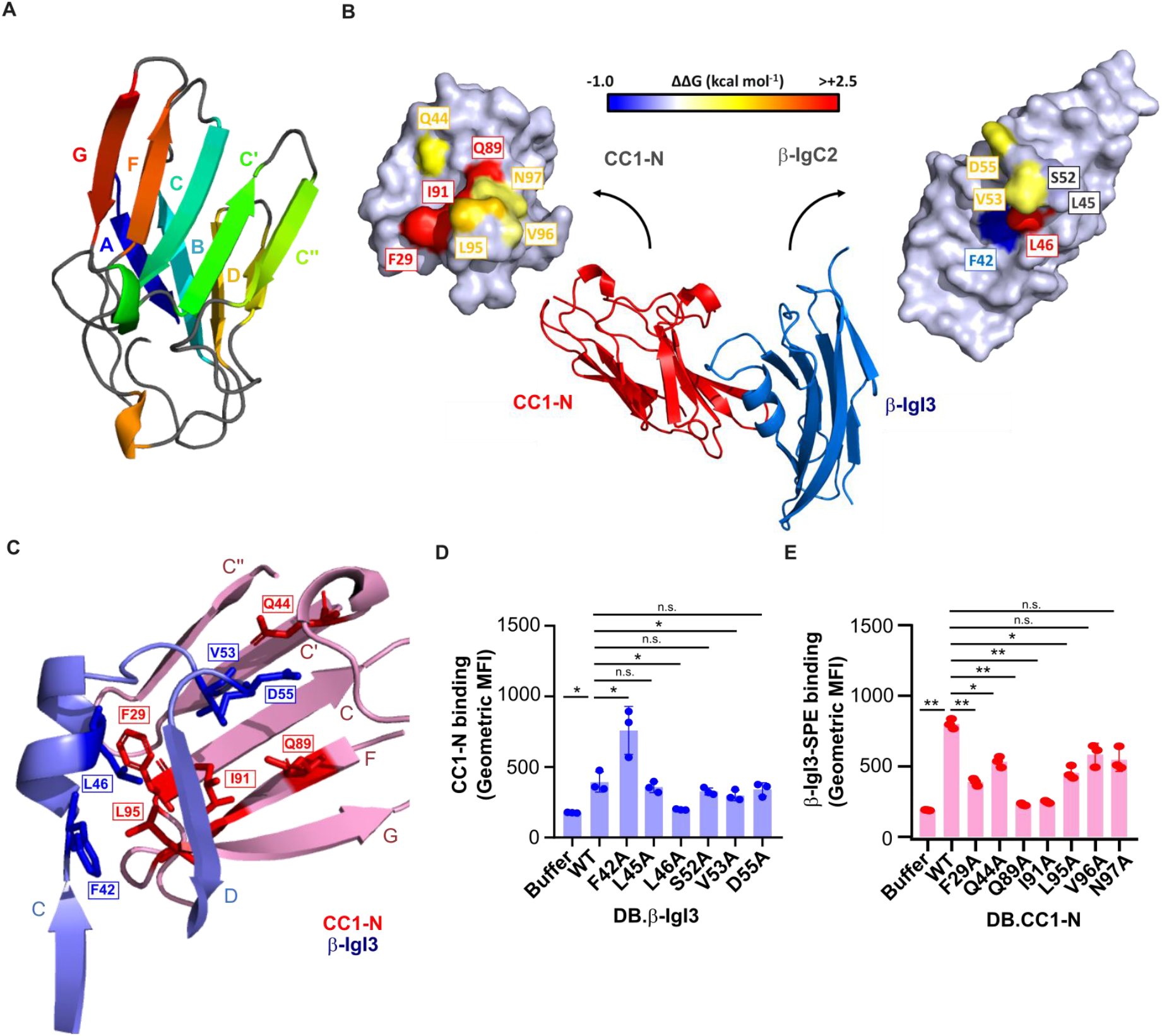
The IgI3 domain of β protein binds CEACAM1 through the unique α-helix and *CD* loop. **A)** Molecular structure of CEACAM1 (CC1)-N, in which each strand of the IgV structure is coloured differenty. **B)** Molecular structure of the β-IgI3 & CC1 N-terminal domain co-complex. Residues that form the β-IgI3-CC1 interface are highlighted. **C)** Close-up of the β-IgI3 (light blue) interface with CC1-N (pink), including highlighting of residues important for forming β-IgI3 (blue) & CC1-N (red) interactions. The *GFCC’C”* strands of CC1-N are shown and labelled, whilst the *C* to *D* strands of β-IgI3 are shown and labelled. **D)** Binding of rCC1-N to dynabeads (DB) coated with β-IgI3 wild type (WT) or mutants. Mutation of β-IgI3 residues 42, 46 and 53 lead to significant changes in CC1-N binding capacity. **E)** Binding of β-IgI3 coupled to Streptavidin-PE to DB coated with rCC1-N wildtype or variants. Mutation of CC1-N residues 29, 44, 89, 91 and 95 lead to significant changes in β-IgI3 binding capacity. Mean and SD values in D and E are displayed for *n* = 3 independent replicates. Statistical significance was calculated using *t* test for D and E, where *, p<0.05; **, p<0.01.

Based on the co-crystal data, we hypothesised that β-IgI3 residue L46 was critical for contacting CEACAM1-N via F29 and L95 (Fig. 6C, Supplementary Fig. 6A & 6B). Additionally, β-IgI3 residue V53 appeared critical for contacting CEACAM1-N residue I91, and β-IgI3 residue F42 was critical for contacting CEACAM1-N residue L95. We generated alanine mutations in β-IgI3 at these positions as well as at several other sites that contact CEACAM1. ITC binding studies of these β-IgI3 mutants to ‘wild-type’ unglycosylated CEACAM1-N reveal that the mutant β-IgI3^L46A^ failed to bind to rCEACAM1-N. Three additional mutants, β-IgI3^L52A^, β-IgI3^V53A^ and β-IgI3^D55A^, had reduced affinity to bind rCEACAM1-N (*K*_D_ = 234±16, 562±44, 690±52 nM, respectively) (Supplementary Fig. 7A; Supplementary Table 6). In contrast, β-IgI3^F42A^ bound with higher affinity (*K*_D_ = 16±15 nM). In addition, we tested the ability of unglycosylated rCEACAM1-N to bind DB coated with the β-IgI3 variants (Fig. 6D). In this analysis, rCEACAM1-N displayed significantly reduced binding to DB coated with β-IgI3^L46A^ and β-IgI3^V53A^, and significantly enhanced binding to DB coated with β-IgI3^F42A^. Together, these data indicate that the L46, L52, V53 and D55 residues of β-IgI3 are critical for the binding to CEACAM1. These residues were observed to be conserved in 57 β protein sequences (data not shown).

For CEACAM1-N, the crystal structure of the complex indicated that β-IgI3 interacts with the dimer interface, contacting several residues which are important for CEACAM1 homodimerization and for binding to other bacterial adhesins. We hypothesised that residues F29, I91 and L95 of CEACAM1 are critical for contacting β-IgI3 (Fig. 6C, Supplementary Fig. 6A & 6B), and generated alanine mutants at these and several other positions in the homodimerization interface. In ITC analysis with unglycosylated CEACAM1-N proteins, both CEACAM1-N^F29A^ and CEACAM1-N^I91A^ failed to bind β-IgI3 (Supplementary Fig. 7B; Supplementary Table 6). Additionally, CEACAM1-N^Q89A^ also lacked ability to bind β-IgI3. Two mutants, CEACAM1-N^Q44A^ and CEACAM1-N^L95A^, resulted in a 10-fold decrease in binding affinities (*K*_D_ = 996±116 and 1350±460 nM, respectively) whilst two further mutants, CEACAM1-N^V96A^ and CEACAM1-N^N97A^, displayed only a modest decrease (4-fold) in binding affinities (*K*_D_ = 370±4 and 490±120 nM, respectively). In addition, we tested the ability of β-IgI3 coupled to streptavidin to interact with the unglycosylated wildtype and mutants rCEACAM1-N-coated DB (Fig. 6E). Binding was significantly reduced for CEACAM1-N^F29A^, CEACAM1-N^Q89A^ and CEACAM1-N^I91A^, as well as for CEACAM1-N^Q44A^ and CEACAM1-N^L95A^, confirming that residues F29, Q89 and I91 are major targets of β-IgI3 binding. I91 has also been identified as a critical CEACAM1 residue for interaction with *M. catarrhalis* (Conners *et al*, 2008), *Neisseria spp*. (Villullas *et al*, 2007; Virji *et al*, 1999), *H. influenzae* (Hill *et al*, 2001) and *Fusobacterium spp* (Wang, 2013). Though Q89 in CEACAM1 is critical for interaction with β-IgI3 and other bacterial ligands, Q89 was not in close contact with any β-IgI3 residues. It is possible that mutation of CEACAM1 residue Q89 forms a gap that I91 attempts to fill that subsequently prevents its interaction with in β-IgI3 residue V53. In summary, contact of residue L46 in β-IgI3 with CEACAM1 residue F29, residue V53 in β-IgI3 with CEACAM1 residue I91 provide the critical interactions. Additional stability is gained through β-IgI3 residues F42 and D55.

### Comparison of β-IgI3- & HopQ-bound CEACAM1 structures

Comparison with the HopQ-CEACAM1 complex, the first shown structure of a bacterial adhesin bound to CEACAM1, revealed that the same set of residues involved in dimerization of CEACAM1 are contacted by both β-IgI3 and HopQ (Supplementary Fig. 6C). However, HopQ is structurally completely unrelated to that of β-IgI3 (Bonsor *et al*, 2018). HopQ uses an intrinsically disordered loop, that folds into a β hairpin and a small helix upon binding to CEACAM1. The β hairpin extends the CEACAM1 *A’GFCC’C”* face whilst the small helix straddles the face (Supplementary Fig. 6C). In contrast, β–IgI3 interacts through the small α helix and loop, with a single residue, L46, critical for β-IgI3 to fit into a pocket on CEACAM1-N formed by F29 and I91 (Fig. 6C). Mutation of any of these residues in β-IgI3 or CEACAM1-N causes a hole in the protein-protein interface or collapse of the pocket, respectively, abolishing binding as observed with our ITC experiments.

### β-IgI3 homologs are broadly distributed in human Gram-positive bacteria

To gain a broader perspective on the possible role of the unique β-IgI3 structure in bacteria-host interaction, we investigated the distribution of this domain in the bacterial kingdom through BLAST analysis. A sequence similar or related to β-IgI3 was identified in 296 proteins predicted to be expressed by human Gram-positive bacteria, including several human pathogens, such are *S. pyogenes, S. dysgalactiae* and *S. pneumoniae*. These sequences formed several different clades upon Maximum Likelihood analysis (Fig. 7A). The β-IgI3 domain was present in a large clade, IgI3 clade I, which also included sequences from *Streptococcus oralis*. A second major clade, clade II, contained sequences from other human streptococcal pathogens including GBS, *S. pyogenes, S. dysgalactiae* and *S. mitis*. Additionally, sequences closely related to clade II were detected in *S. mitis, S. milleri* and *S. intermedius*, forming clade III. Finally, many homologs of β-IgI3 were present in proteins from a diverse panel of Gram-positive human pathogens including *S. pneumoniae, S. pseudopneumoniae*, and *Gemella haemolysans*. In addition, a homolog was identified in *Gardnerella vaginalis*, a Gram-variable bacterium. The sequences of β-IgI3 homologs were all located in proteins predicted to be surface-localized and anchored in the bacterial cell wall.

**Figure 7:**
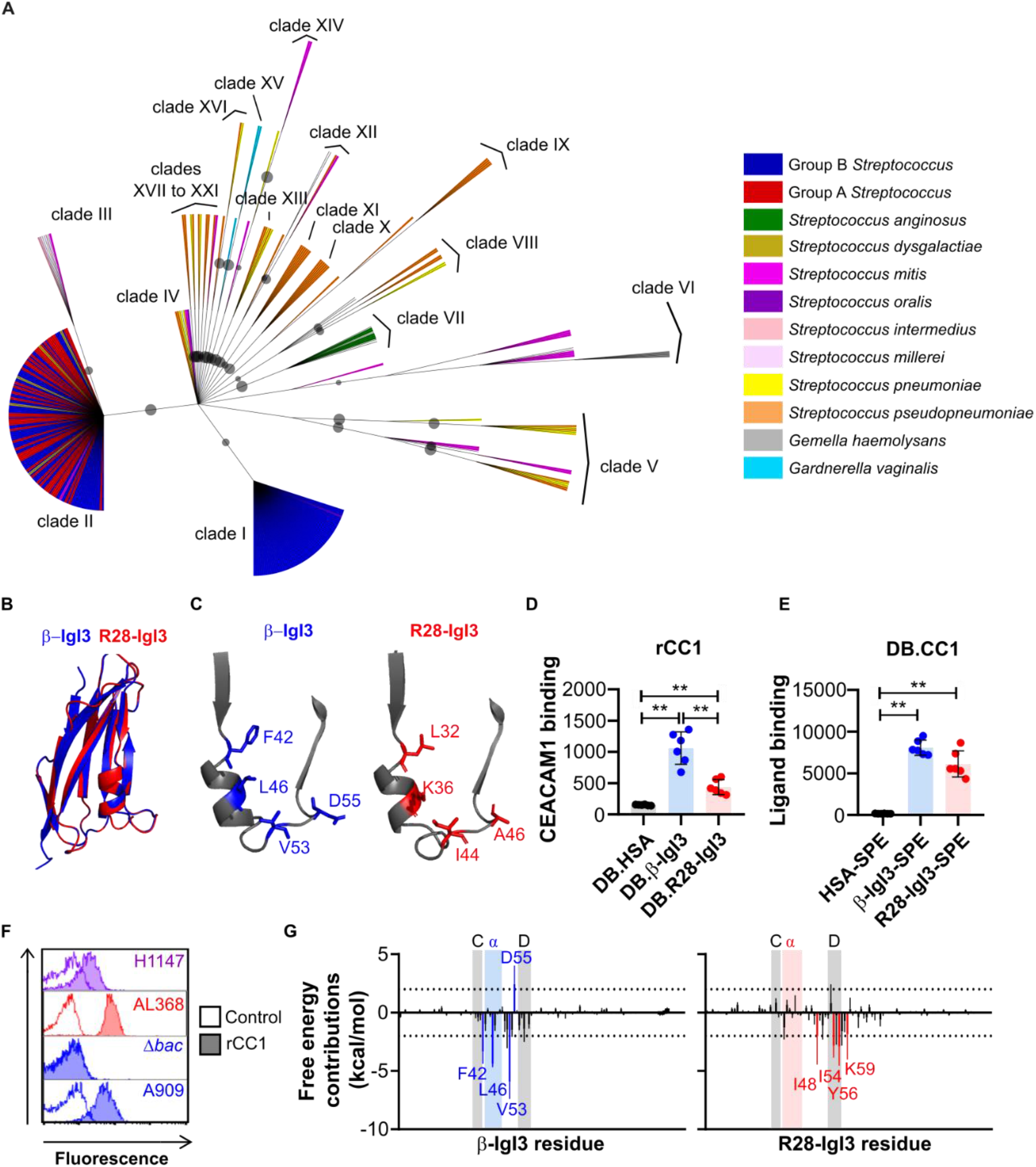
The unique IgI3 structure is widely present in Gram-positive bacterial species that bind human CEACAM1. **A)** Amino acid based Maximum Likelihood tree of β-IgI3 homologs. Clades of related IgI3 domains are shown, with clade I representative of β-IgI3 domain sequences. Branches are coloured by bacterial species from which the sequence originates, except those with no species designated. Maximum likelihood values bootstrap values > 60 are shown as a circle, where larger circles indicate a higher bootstrap support value. **B)** Superposition of the IgI3 domain from β protein (blue) onto predicted IgI3 domain structure of clade II sequence of *S. pyogenes* R28 protein (red). **C)** Close-up of the truncated *C* strand, α-helix and *D* strand of β-IgI3 (blue) and R28-IgI3 (red), indicating the corresponding R28-IgI3 residues present at sites in β-IgI3 that bind CEACAM1 (CC1). **D)** Binding of recombinant (r)CC1-HIS to dynabeads (DB) coated with β-IgI3, R28-IgI3 or control. Mean and SD is shown for *n* = 6 independent replicates. **E)** Binding of β-IgI3, R28-IgI3 or human serum albumin (HSA), coupled to PE-conjugated streptavidin (SPE), to DB coated with CC1 (DB.CC1). Mean and SD is shown for *n* = 6 independent replicates. **F)** The per residue free energy contribution spectrums of β-IgI3/CC1-N interface and R28-IgI3/CC1-N interface. The residues with binding free energy contributions lower than −2.0 kcal/mol are identified as key (favourable) residues and coloured blue (for β-IgI3) or red (for R28-IgI3). Free energy contributions at −2.0 kcal/mol and 2.0 kcal/mol are indicated by dashed lines. The position of the truncated *C* strand, α-helix and *D* strand is shaded for each protein. The mean and standard deviation is representative of estimates from 50 docked models. **G)** Binding of recombinant rCC1-HIS (10 μg/mL) to R28-expressing GBS strain H1147 and R28-expressing *S. pyogenes* strain AL368. Data representative of *n* = 3 replicates. Statistical significance was calculated using a One-way ANOVA with Sidak’s multiple comparisons for D and E, where *, p<0.05; **, p<0.01.

We predicted the tertiary structure of 11 representative homolog sequences based on the β-IgI3 crystal structure. These structures maintained the overall IgI3 fold, including an α-helix and loop located between the truncated *C* strand and the *D* strand (Supplementary Fig. 8), except clade XV sequence from *G. vaginalis* that lacked the *C* strand. These data suggest that the IgI3 structure is broadly distributed in bacterial cell wall-anchored adhesins? We focused further analysis on clade II that maintains key IgI3 structures despite sharing only 40% amino acid identity with β-IgI3 (Fig 7B; Supplementary Fig. 9A, 9B & 9C). This clade includes the R28 protein, which is found in *S. pyogenes* and in GBS (where the same protein is also called Alp3). R28 is a cell wall-anchored adhesin that is unrelated to β protein except for the IgI3 domain (Stålhammar-Carlemalm *et al*, 1999; Lindahl *et al*, 2005). To understand the CEACAM1 binding capacity of β-IgI3 homologs, we inspected whether the critical protruding residues located between *C* and *D* strands in β-IgI3 (Fig. 5G) were conserved across the homologs. Though the CEACAM1-binding residues of β-IgI3 (F42, L46, V53 and D55) are not conserved in R28-IgI3 (Fig. 7C), two of the R28-IgI3 residues found at corresponding positions were of hydrophobic nature as in β-IgI3. Interestingly, additional residue variation was evident among other IgI3 domains (Supplementary Fig. 11). Since this result suggested that IgI3 homologs are likely to differ in their properties, it became critical to analyze whether the divergent β-IgI3 homolog in the R28 protein binds CEACAM1.

### Human CEACAMs bind Gram-positive bacteria expressing β-IgI3 homologs

To determine whether the IgI3 domain in R28 interacts with CEACAM1, we purified rR28-IgI3 protein domain from *E. coli* and tested interaction with rCEACAM1. Beads coated with biotinylated R28-IgI3 and β-IgI3, but not HSA, interacted with rCEACAM1 (Fig. 7D). In the reverse assay, beads coated with rCEACAM1 interacted with R28-IgI3 and β-IgI3, but not HSA, coupled to streptavidin (Fig. 7E). We also confirmed that rCEACAM1 bound to an R28-expressing *S. pyogenes* strain (Fig. 7F). In addition, rCEACAM1 bound to an Alp3-expressing strain of GBS, which was expected given that R28 and Alp3 proteins are identical in sequence. Taken together with the structural predictions described above, these data indicate that IgI3 folds from a wider range of Gram-positive bacterial pathogens can interact with human CEACAM1, but individual IgI3 domains may interact with human CEACAM1 differently.

To dissect whether β-IgI3 and R28-IgI3 interact differently with CEACAM1, we simulated the docking of IgI3 domains onto human CEACAM1 and examined the free energy calculations to ascertain the key IgI3 residues in the complex formation (Supplementary Fig. 9C). Residues with binding free energy contributions lower than −2.0 kcal/mol and greater than 2 kcal/mol are identified as key residues and unfavourable residues, respectively. Simulation of β-IgI3 and CEACAM1 docking correctly predicted residues F42, L46 and V53 were key determinants of CEACAM1 binding, whilst D55 was an unfavourable determinant (Fig 7G; Supplementary Fig. 9D). The binding free energy of the (R28-IgI3)-(CEACAM1-N) complexes was strong (−31.02±6.70 kcal/mol, n = 50 simulations). In contrast to β-IgI3, residues located in the α-helix of R28-IgI3 were not identified as key determinants of CEACAM1 binding (Fig 7G; Supplementary Fig. 9E). Instead, critical R28-IgI3 residues (K48, I54, I56 and K59) were located in the *CD* loop and the *D* strand only. Mapping of key residues onto the surface structure suggest that β-IgI3 and R28-IgI3 target CEACAM1 through different faces on the IgI3 fold (Supplementary Fig. 9D and 9E). Thus, IgI3 domains in different bacterial proteins probably interact with human CEACAM1 through mechanisms that are not identical but use the same structural fold.

## Discussion

We show here that a novel Ig-like fold in different streptococcal adhesins targets human CEACAM receptors. Molecular analysis of this domain in β protein from GBS demonstrated that the Ig-fold binds to the IgV-like N-terminal domains of human CEACAM1 and CEACAM5. Additional biochemical analysis demonstrated that the interaction is of high specificity and high-affinity. We solved the crystal structure of the Ig-like fold in β and revealed it represents a novel Ig-fold structure related to the I-set. The Ig-like fold identified in β is characterized by an absence of cysteine residues and the presence of a truncated *C* strand that is directly followed by a unique 1.5-turn α-helix. As this Ig-like fold of the I-set has unique features, it was designated IgI3. The absence of cysteine residues has been reported in Ig folds,(Halaby & Mornon, 1998) but has not been documented in IgI folds to date. In addition, the partial replacement of the *C* strand with an 1.5-turm α-helix was not identified in any other Ig fold in a search of PDB using HHpred (Zimmermann *et al*, 2018). I-set folds are rarely found within bacterial proteins as observed through a search of the SMART database (Letunic & Bork, 2018). This work presents the first representative structure of a novel Ig-fold subtype and highlights that unique subtypes of Ig folds remain uncharacterised in biology.

The co-crystal structure of the complex demonstrated that β-IgI3 targets the dimerization interface of CEACAM1 through a unique mechanism. In particular, the streptococcal β protein uses the unique features of the IgI3 domain to bind the CEACAM homodimerization face. Specifically, the 1.5-turn α-helix in β-IgI3 interacts with the *C* and *F* strands of CEACAM1 using residue L46 for binding, whilst the β-IgI3 *CD* loop is also involved in critical interaction with the *F* strand. Therefore, the unique structural features of the IgI3 domain are responsible for forming CEACAM1 interactions. In contrast, Opa proteins of *Neisseria spp* are membrane proteins that bind to the CEACAM dimerization interface through two hypervariable loops (Billker *et al*, 2000). UspA1 of *M. catarrhalis* forms a trimeric coiled-coil that binds to CEACAM dimerization interface through a small bend (Conners *et al*, 2008), HopQ of *H. pylori* uses a disordered loop to bind the CEACAM dimerization interface (Bonsor *et al*, 2018). and AfaE of *E. coli* uses an incomplete Ig-like domain to target the CEACAM dimerization interface (Anderson *et al*, 2004; Korotkova *et al*, 2008).

The finding that β-IgI3 homologs are broadly distributed in Gram-positive bacteria raises many fascinating questions about the evolutionary history of the IgI3-fold and their role in bacterial pathogenesis. What is the origin of the IgI3-fold? How conserved is the IgI3-fold structure? How did the IgI3-fold become so widely distributed? Do all bacterial adhesins containing IgI3 homologs bind CEACAMs and how is binding conferred? IgI3 homologs were found in adhesins from several Gram-positive bacteria. Structural modelling of representative sequences predicted that the homologs have a structure resembling β-IgI3. There was low conservation of the critical β-IgI3 residues, suggesting that each sequence may possess different properties. Our study of one homologous domain with divergent sequence, R28-IgI3 in clade II, demonstrated that it can bind to CEACAMs. As R28-IgI3-like sequences were found in proteins from multiple human pathogens, it is tempting to speculate that CEACAM binding may represent an unrecognized adhesion strategy for a range of Gram-positive bacteria. Thus, our data reveals that Gram-positive bacteria, in addition to Gram-negative bacteria, have responded to the strong selective pressure to evolve CEACAM-binding adhesins (Tchoupa *et al*, 2014). Of note, the genes encoding β and R28 in GBS are encoded on mobile genetic elements that can move between bacteria by horizontal gene transfer (Tettelin *et al*, 2005; ben Zakour *et al*, 2012). This could provide opportunities for rapid acquisition of CEACAM-binding in other bacterial species leading to the emergence of strains with enhanced virulence. Intriguingly, simulated docking suggested that R28-IgI3 and β-IgI3 bind through alternative strategies. Thus, the exploration of IgI3-fold and CEACAM receptor interactions is worthy of further exploration.

Our data here show that GBS can bind CEACAMs through expression of IgI3 -fold containing adhesins. This is of importance as there is little knowledge of the cellular adhesion mechanisms utilized by GBS, despite its role as a major neonatal pathogen. CEACAM engagement has also independently evolved in several Gram-negative human pathogens (Tchoupa *et al*, 2014, 2015; Königer *et al*, 2016; Brewer *et al*, 2019; Conners *et al*, 2008; Virji *et al*, 1996; Hill *et al*, 2001; Bos *et al*, 1998). It is likely that β-expressing GBS also adhere to human cells via CEACAMs, similarly to the Gram-negative bacteria (Bos *et al*, 1998; Gutbier *et al*, 2015; Tchoupa *et al*, 2014, 2015; Königer *et al*, 2016; Javaheri *et al*, 2016; Virji *et al*, 1996). Our *in vitro* adhesion assays assessing the β-dependent mechanism support this conclusion. Thus, CEACAM1 and CEACAM5 expand the repertoire of human cell surface proteins that GBS may employ for adhesion (Pietrocola *et al*, 2018). Further work is required to determine the consequences of the interaction between β-IgI3 and CEACAM beyond cell lines.

The β protein of GBS is commonly expressed by serotype Ia, Ib, II and V strains (Lindahl *et al*, 2005). It was recently shown that high β protein expression levels are associated with increased virulence of GBS clinical isolates.(Nagano *et al*, 2006) As β protein binds the inhibitory receptor CEACAM1, as well as the inhibitory receptors Siglec-5 and −7 that are expressed on leukocytes, we speculate that dual- or multi-engagement of these inhibitory receptors may induce potent immunosuppressive signals through induction of tyrosine phosphorylation (Carlin *et al*, 2009; Fong *et al*, 2018). Moreover, the ability of the β protein to bind other components of the human immune system, *viz*. the plasma proteins IgA and factor H (Lindahl *et al*, 1990; Russell-Jones *et al*, 1984; Hedén *et al*, 1991), could enable it to act as a multitool protein that assists epithelial cell adhesion, suppresses immune cell activity and promotes immune evasion. A similar multifunctional protein is UspA1 of *M. catharralis*, which interacts with CEACAMs, fibronectin, complement factor C3 and the complement regulator C4BP (Lindahl *et al*, 1990; Tong Tan *et al*, 2005; Nordström *et al*, 2005). Future studies are warranted to improve our knowledge of the role of β protein as a virulence factor.

In summary, our data demonstrate that an adhesin containing a domain with unique Ig-like fold, the IgI3 fold, promotes binding of GBS to human CEACAM receptors. Moreover, homologous domains were identified in a variety of Gram-positive bacterial including pathogens, suggesting that many Gram-positive bacterial pathogens employ IgI3 domains to engage human CEACAMs. Characterisation of these domains may lead to a better understanding of pathogenicity mechanisms and could pave-the-way for the development of novel therapeutic approaches to prevent infections.

## Methods

### Bacteria growth conditions

Organisms used in this study are listed in Supplementary Table 1. Bacteria were cultured in Todd-Hewitt Broth (TH; *S. agalactiae*), TH Broth + 1% yeast (*S. pyogenes*), Tryptic Soy Broth (TSB; *S. aureus*) or Lysogeny Broth (LB; *E. coli*) and supplemented with antibiotics as listed in Supplementary Table 1.

### Expression and purification of glycosylated and unglycosylated rCEACAMs

Glycosylated CEACAMs were expressed and purified from Expi29F cells (Life Technologies). In brief, gBlocks containing open reading frames (ORFs) encoding the extracellular domain of CEACAMs, and a C terminal LPETGGSHHHHHH tag, were cloned into the pcDNA3.4 expression vector (Invitrogen) (Supplementary Table 2). Recombinant CEACAM-His proteins were expressed in Expi293F cells (Life Technologies) and purified by affinity chromatography (ÄKTA Pure, GE Healthcare Life Sciences) using a Nickel column (GE Healthcare Life Sciences) as previously described (Zhao Y *et al*, 2020). Eluate was dialysed against 300 mM NaCl 50 mM Tris pH 7.8 at 4°C. Unglycosylated CEACAMs were expressed and purified from *E. coli* cultures. In brief, gBlocks containing ORFs encoding the N-terminal or A1B1A2 domains of CEACAM1, and a C terminal LPETGGS-6xHis tag, were cloned into the pRSET-C vector (Supplementary Table 2). Vectors were transformed into the Rosetta Gami (RG) *E. coli* strain. RG strains were cultured in LB supplemented with 100 μg/ml ampicillin and 1 mM D-glucose at 37°C overnight, and subcultured 1:20 into LB supplemented with 1 mM D-glucose at 37°C until OD_600_ = 0.4. Protein expression was induced by culturing for 4 hours at 37°C following the addition of 1 mM IPTG. After lysis of bacteria, proteins were purified using a Nickel column (GE Healthcare Life Sciences) and affinity chromatography (ÄKTA Pure, GE Healthcare Life Sciences). Eluates dialysed against 300 mM NaCl 50 mM Tris pH 7.8 at 4°C. Large scale expression of tag-less CEACAM1-N domain and point mutants for use in isothermal titration calorimetry (ITC) and crystallization were expressed using pET21d vectors in *E. coli* and purified as previously described (Bonsor *et al*, 2015a).

### Expression and purification of bacterial proteins

β, α and Rib proteins were purified from GBS cultures as previously described (Lindahl *et al*, 1990; Stålhammar-Carlemalm *et al*, 1993). Domains of the β proteins (B6N, IgABR, B6C, IgSF and β75KN) and R28 (R28-IgI3) were cloned and expressed in *E. coli* (Supplementary Table 2). The IgSF domain structure was found to be in a region overlapping the previously published B6C (amino acids 226-434) and β75KN domains (amino acids 434-788). (Lindahl *et al*, 2005; Areschoug *et al*, 2002b) Therefore, we have updated domain designations as follows:-B6C represents amino acids 226-390, IgSF represents amino acids 391-503, and β75KN represents amino acids 504-788. Briefly, gBlocks containing ORFs and a C-terminal LPETGGS-6xHis tag, were cloned into the pRSET-C vector using BamHI and NdeI. Vectors were transformed into the RG strain. RG strains were cultured, induced, harvested and lysed as described above for unglycosylated CEACAMs. Bacterial supernatants were filtered, supplemented with 10 mM imidazole and passed over a Nickel column (GE Healthcare Life Sciences). Proteins were purified by affinity chromatography (ÄKTA Pure, GE Healthcare Life Sciences), eluted with imidazole and were dialysed against 300 mM NaCl 50 mM Tris pH 7.8 at 4°C.

### Binding of glycosylated and non-glycosylated recombinant CEACAM to bacteria

Six x 10^6^ of mid-logarithmic phase bacteria were incubated with rCEACAM (1, 3, 5, 6 or 8), rCEACAM1-N or rCEACAM1-A1B1A2 at 4°C for 1 hour with shaking. After washing in PBS + 0.1% bovine serum albumin (BSA), rCEACAM on the bacterial surface was detected by incubation with FITC-conjugated anti-HIS monoclonal antibody (mAb) at 4°C for 1 hour with shaking. Bacterial fluorescence was measured by flow cytometric analysis following washing and fixation in 1% paraformaldehyde (PFA). To detect rCEACAM binding to GBS using pull-down Western blot analysis, bacterial pellets were resuspended in 1x sample buffer and heated to 95°C for 10 mins. Lysates were separated by SDS-PAGE in a 12.5% polyacrylamide gel at 270V, blotted onto nitrocellulose membranes and probed with anti-CEACAM1 (clone C51X/8) mAb. Membranes were probed using rabbit anti-mouse-IgG-HRP and developed using ECL substrate.

### Measurement of β protein expression by GBS strains

Rabbit antiserum was raised against β protein as previously described.(Lindahl *et al*, 1990) Six x 10^6^ of mid-logarithmic phase bacteria were incubated with heat-inactivated 0.1% rabbit anti-β serum or normal rabbit serum at 4°C for 1 hour with shaking, washed and incubated in the presence of PE-conjugated goat anti-rabbit-IgG at 4°C for 1 hour with shaking. Fluorescence of bacteria was measured by flow cytometric analysis after washing and fixation in 1% PFA.

### Detection of CEACAM1 and β protein interaction

#### Western blot analysis

Purified GBS proteins, or β protein domains (B6N, IgABR, B6C, IgSF or β75KN), were separated by non-reducing SDS-PAGE, transferred to nitrocellulose membranes, blocked with 4% non-fat dried milk overnight at 4°C, probed with 10 μg/mL rCEACAM1-His (4°C) and horse radish peroxidase (HRP)-conjugated mouse anti-His-IgG, and developed using Pierce ECL Western Blot Substrate Reagent (ThermoFisher Scientific).

#### Binding of bacterial proteins to CEACAM1-coated dynabeads

rCEACAM-His was attached to nickel dynabeads (DB; Dynabead His-tag pull-down & isolation, Invitrogen) following standard protocols. Presence of CEACAM1 was confirmed by incubating DB in the presence of 5 μg/mL anti-CEACAM or isotype control mAb, and detection with FITC-conjugated mouse anti-His-IgG mAb. To detect binding of purified GBS proteins, 40 μg of DB was incubated in the presence of 10 μg/mL of β protein and probed with 0.1% mouse antiserum or normal mouse serum, and secondary goat anti-mouse-IgG-PE mAb. Biotinylated proteins (including β-IgSF-biotin or HSA-biotin amongst others) were incubated with PE-conjugated Streptavidin on ice for 10 mins. This generates biotin-streptavidin complexes on the remaining 3 binding pockets of streptavidin. To detect binding of proteins coupled with streptavidin (β-IgSF-biotin or HSA-biotin formed with Streptavidin-PE), 20 μg of DB was incubated with varying concentrations of protein-streptavidin. All dilutions and DB washes employed PBS + 0.1% BSA, all incubations were performed for 1 hour at 4°C, and all fixations used 1% PFA. Fluorescence of DB was measured by flow cytometry. For inhibition experiments, CEACAM-coated DB were pre-incubated with 5 μg/ml anti-CEACAM1-N (clone CC1/3/5-Sab, LeukoCom, Essen, Germany) or anti-CEACAM1-A1B1 mAb (clone B3-17, LeukoCom, Essen, Germany) or 10 μg/mL HopQ for 1 hour at 4°C.

#### Binding of CEACAM to bacterial protein coated dynabeads

Proteins with C-terminal biotin tags were attached to streptavidin-coated dynabeads (Dynabead M-280 Streptavidin, Invitrogen) following standard procedures. To detect binding of rCEACAM-His or rCEACAM-N-His, 3 x 10^5^ DB were incubated in the presence of 9 μL of proteins (concentrations range 0.1 to 30 μg/mL), and probed with FITC-conjugated mouse anti-His-IgG. In the case of rSiglec-5-Fc, binding was detected using anti-IgG-AF488. All dilutions and DB washes employed PBS + 0.1% BSA, all incubations were performed for 1 hour at 4°C, and all fixations used 1% PFA. Fluorescence of DB was measured by flow cytometry.

### Isothermal titration calorimetry

ITC measurements were performed on an iTC200 instrument (GE Healthcare). Typically 500 μM of unglycosylated CEACAM-N domains were loaded into the syringe of the calorimeter and 50 μM β-IgC2 protein was loaded into the syringe. All measurements were performed at 25 °C, with a stirring speed of 750 rpm., in 30 mM Tris-HCl, 150 mM NaCl, pH 7.5. Data were analyzed using the Origin 7.0 software.

### Adhesion of GBS to epithelial cells

HeLa and Chinese Hamster Ovary (CHO) cell lines expressing CEACAMs were cultured as previously described (Bos *et al*, 1998; Hollandsworth *et al*, 2020; Daniel *et al*, 1993), and seeded into 24-well tissue culture plates. Tightly confluent monolayers were infected with mid-logarithmic phase (absorbance at OD_600_ = 0.4) GBS strains at a multiplicity of infection (MOI) of 10. Assays were commenced by centrifugation for 2 minutes at 700 rpm, and incubated at 37°C with 5% CO2 for 30 mins. Cells were washed five times with DMEM, detached with 0.25% trypsin, and lysed (PBS + 0.025% Triton-X). Adherent bacteria were enumerated using serial dilution plating and growth on Todd-Hewitt agar plates. In specific assays, cells were pre-incubated with 5 μg/mL mAb (anti-CEACAM1-N or anti-CEACAM1-A1B mAb) for 60 mins, or bacteria were pre-incubated with rCEACAM1-N for 60 minutes before co-culture. For confocal microscopy, experiments were performed with FITC-labelled GBS at a MOI of 10 for 30 minutes. After washing five times with PBS, monolayers were fixed with 4% PFA overnight prior to staining of nuclei with DaPI (Sigma Aldrich) and cell membranes with AF647-conjugated wheat germ agglutinin (ThermoFisher Scientific). To perform assays with detached CHO cell lines, adherent monolayers were detached by incubation at 37°C for 5 mins with Accutase Cell Detachment Solution (BioLegend). After washing, cells were re-suspended to 3 x 10^6^ cells/ml in DMEM containing 10% FCS. To measure CEACAM expression, cells were incubated with mouse mAb clone CC1/3/5-Sab (detects N domain of human CC1, CC3 and CC5) or 5C8C4 (detects A1B1A2B2A3B3 domain of human CC5; LeukoCom, Essen, Germany) for 30 minutes, washed and incubated with PE-conjugated rabbit anti-mouse-IgG. To measure GBS binding, cells were incubated with FITC-labelled GBS at a MOI of 10 for 30 minutes at 4°C. After washing with PBS, cells were fixed in 4% PFA and fluorescence was measured by flow cytometry.

### Crystallography of unglycosylated CEACAM1-N and β-IgSF

Initially, the β-IgSF & CEACAM1-N complex was prepared by mixing the proteins with a slight excess of CEACAM1-N (1:1.1 ratio). Protein was concentrated using an amicon Ultra-4 10,000 NMWL before purification by size exclusion chromatography using a Superdex 200 column (GE Healthcare) equilibrated with 20 mM Tris, 150 mM NaCl, pH 7.5. The complex was concentrated to 9 mg/mL and screened against the commercial Wizards I/II/III/IV (Rigaku) and JCSG^+^ (QIAgen) screens using a Crystal Gryphon Protein Crystallography System (Art Robbins Instruments). Crystals were found in several conditions but diffraction was never found beyond 8.0 Å in the best condition of 1.8 M Ammonium Sulfate, 0.1 M Sodium Citrate pH 5.5. β-IgSF contains both a C-terminal Sortase and 6xHis tag. This was removed by first incubating β-IgSF by itself with Carboxypeptidase A (Sigma). 2 mL of 6 mg mL^−1^ of β-IgSF was incubated with 1:100 w/w of Carboxypeptidase A for 24 hours at 4 °C before the addition of another 1:100 w/w of Carboxypeptidase A and incubation for 24 hours at room temperature. The reaction was terminated by the addition of EDTA to a final concentration of 5 mM. The protein was concentrated using an Amicon Ultra-4 10,000 NMWL before purification by size exclusion chromatography using a Superdex 200 column (GE Healthcare) equilibrated with 20 mM Tris, 150 mM NaCl, pH 7.5. CEACAM1-N was added in a slight excess (1:1.05 ratio) and purified by size exclusion as described above. The complex was concentrated and screened again at a concentration of 7.5 mg mL^−1^. Crystals again formed in the condition of 1.8 M Ammonium Sulfate, 0.1 M Sodium Citrate pH 5.5, however their morphology were different. Crystals were cyroprotected in 30 % v/v glycerol. Anisotropic diffraction was typically observed with diffraction to ~3.0 Å in two directions and 4.0 Å in the other. Two datasets were collected; (i) beamline ID23-D at the Advance Photon Source and (ii) beamline 12-2 at the Stanford Synchrotron Radiation Lightsource. Data was processed and indexed by XDS, merged together with BLEND before scaling and conversion to structure factors using Aimless. Matthew’s coefficient suggests five complexes in the asymmetric unit. Molecular replacement using MOLREP and CEACAM1-N (PDB entry 2gk2) as a search model in MOLREP only found two CEACAM1 molecules. Two copies of the complex would suggest a high solvent content (78 %). Refinement using REFMAC5 and visualization in Coot, showed that CEACAM1-N was well placed and residual electron density was present for the β-IgI3 protein. Density modification was performed using Parrot (solvent flattening, histogram matching and NCS averaging) to improve the maps. The β-IgSF backbone was built manually. A PDBeFold search of the backbone showed that it was an IgI-like fold, with the closest fold by R.M.S.D being the IgC2 domain of Perlecan (PDB entry 1gl4). The Ig fold of β-IgSF possesses a previous unrecognized modification (Fig 5B and 5C). We denote this as the I3-set domain, and the domain with β protein as β-IgI3. This was used to successfully position sidechains, lock the registry and provide extra restraints during refinement in REFMAC5 with ProSMART. The (β-IgI3)-(CEACAM1-N) complex has been deposited to the PDB with the entry code, 6V3P. Atom contacts in the (β-IgI3)-(CEACAM1-N) complex interface were identified using NCONT (CCP4) with a cut-off of 4.0Å (Winn *et al*, 2011).

### Bioinformatics analysis

BLAST analysis of the β-IgI3 amino acid sequence was performed to identify homologs in bacterial proteins, and subsequently aligned using ClustalW. Phylogenetic analysis was performed using Maximum Likelihood approach and 1000 bootstrap replications in MEGA (Kumar *et al*, 2016), and the resulting tree displayed with interactive tree of life version 4 (Letunic & Bork, 2019). The structure of IgI3 homologs was predicted using the β-IgI3 structure as input in SWISS-MODEL (Webb & Sali, 2016). Only models passing a QMEAN score of < −4.00 were further analyzed. Docking of β-IgI3 structure or the R28-IgI3 predicted structure to CEACAM1-N was simulated 50 times using ZDOCK server (Pierce *et al*, 2014), in which CEACAM1-N residues F29, Q44, Q89, I91 and N95 were included as contact sites. The free energy contribution of each simulation was interpedently calculated using MM/GBSA analysis on the HAWKDOCK server (Weng *et al*, 2019).

### Data Availability

The co-crystal structure of β-IgI3 and CEACAM1-N is available at the PDB with ID: 6V3P. All other data supporting the findings of this study are available within the paper and its supplementary information files and are available from the corresponding author on reasonable request.

## Acknowledgments

The authors thank Carla J.C. de Haas, Piet Aerts, Kok P.M. van Kessel (UMC Utrecht, Utrecht, The Netherlands), Birgit Maranca-Hüwel, Bärbel Gobs-Hevelke (University of Duisberg-Essen, Germany) and Margaretha Stålhammar-Carlemalm (Lund University, Sweden) for excellent technical support and invaluable help. We thank Paul Sullam (University of California, USA) for providing *S. sanguinis* strains. We also wish to thank the support staff of Beamlines of 12-2 and ID23-D at the Stanford Synchrotron Radiation Lightsource and the Advanced Photon Source for their aid in data collection, respectively. We thank Irene M. Mavridis (Institute of Child Health, Greece) and José R. Penadés (Imperial College London, UK) for critical reading of the manuscript. This work was supported by the European Union’s Horizon 2020 Research and Innovation Programme under Grant Agreement 700862, Deutsche Forschungsgemeinschaft DFG grant SI-1558/3-1, The Swedish Research Council, The Foundation Olle Engkvist Byggmästare, and the National Institutes of Health R01 NS116716 (to K.S.D.)

## Author Contributions

Conception & design of the work (N.M.V., L.D., D.A.B., E.J.S., J.A.G.V., K.S.D., B.B.S., G.L., A.J.M.). Acquisition, analysis or interpretation of data (N.M.V., L.D., D.A.B., J.B., V.S., M.L., A.S., O.R.N., E.B., E.L., J.A.G.V., K.S.D., B.B.S., G.L., A.J.M.). Drafting of manuscript (N.M.V., L.D., D.A.B., J.A.G.V., K.S.D., B.B.S., G.L., A.J.M.). All authors read and commented on the final version of the manuscript.

## References

Ali SR, Fong JJ, Carlin AF, Busch TD, Linden R, Angata T, Areschoug T, Parast M, Varki N, Murray J, Nizet V & Varki A (2014) Siglec-5 and Siglec-14 are polymorphic paired receptors that modulate neutrophil and amnion signaling responses to group B Streptococcus. Journal of Experimental Medicine 211: 1231–1242

Anderson KL, Billington J, Pettigrew D, Cota E, Simpson P, Roversi P, Chen HA, Urvil P, du Merle L & Barlow PN (2004) An Atomic Resolution Model for Assembly, Architecture, and Function of the Dr Adhesins. Molecular Cell 15: 647–657

Areschoug T, Linse S, Stålhammar-Carlemalm M, Hedén L-O & Lindahl G (2002a) A Proline-Rich Region with a Highly Periodic Sequence in Streptococcal Protein Adopts the Polyproline II Structure and Is Exposed on the Bacterial Surface. Journal of Bacteriology 184: 6376–6383

Areschoug T, Stålhammar-Carlemalm M, Karlsson I & Lindahl G (2002b) Streptococcal β protein has separate binding sites for human factor H and IgA-Fc. Journal of Biological Chemistry 277: 12642–12648

Arredondo-Alonso S, Top J, Mcnally A, Puranen S, Pesonen M, Pensar J, Marttinen P, Braat JC, Rogers MRC, van Schaik W, Kaski S, Willems RJL, Corander J, Schürch AC & Projan SJ (2020) Plasmids Shaped the Recent Emergence of the Major Nosocomial Pathogen Enterococcus faecium Downloaded from. mBio 11: e03284–19

Baba T, Bae T, Schneewind O, Takeuchi F & Hiramatsu K (2008) Genome sequence of Staphylococcus aureus strain newman and comparative analysis of staphylococcal genomes: Polymorphism and evolution of two major pathogenicity islands. Journal of Bacteriology 190: 300–31

Banerjee A, Kim BJ, Carmona EM, Cutting AS, Gurney MA, Carlos C, Feuer R, Prasadarao N v. & Doran KS (2011) Bacterial Pili exploit integrin machinery to promote immune activation and efficient blood-brain barrier penetration. Nature Communications 2: 462

Bateman A, Eddy SR & Chothia C (1996) Members of the immunoglobulin superfamily in bacteria. Protein Science 5: 1

Bensing BA, Loukachevitch L v., Mcculloch KM, Yu H, Vann KR, Wawrzak Z, Anderson S, Chen X, Sullam PM & Iverson TM (2016) Structural basis for sialoglycan binding by the streptococcus sanguinis SrpA adhesin. Journal of Biological Chemistry 291: 7230–7240

Billker O, Popp A, Gray-Owen SD & Meyer TF (2000) The structural basis of CEACAM-receptor targeting by neisserial Opa proteins. Trends in Microbiology 8: 258–260

Bolduc GR & Madoff LC (2007) The group B streptococcal alpha C protein binds α1β1-integrin through a novel KTD motif that promotes internalization of GBS within human epithelial cells. Microbiology 153: 4039–4049

Bonsor DA, Beckett D & Sundberg EJ (2015a) Structure of the N-terminal dimerization domain of CEACAM7. Acta Crystallographica Section:F Structural Biology Communications 71: 1169–1175

Bonsor DA, Günther S, Beadenkopf R, Beckett D & Sundberg EJ (2015b) Diverse oligomeric states of CEACAM IgV domains. Proceedings of the National Academy of Sciences of the United States of America 112: 13561–13566

Bonsor DA, Zhao Q, Schmidinger B, Weiss E, Wang J, Deredge D, Beadenkopf R, Dow B, Fischer W, Beckett D, Wintrode PL, Haas R & Sundberg EJ (2018) The Helicobacter pylori adhesin protein HopQ exploits the dimer interface of human CEACAMs to facilitate translocation of the oncoprotein CagA. The EMBO Journal 37: e98664

Bos MP, Kuroki M, Krop-Watorek A, Hogan D & Belland RJ (1998) CD66 receptor specificity exhibited by neisserial Opa variants is controlled by protein determinants in CD66 N-domains. Proceedings of the National Academy of Sciences of the United States of America 95: 9584–9589

Brewer ML, Dymock D, Brady RL, Singer BB, Virji M & Hill DJ (2019) Fusobacterium spp. target human CEACAM1 via the trimeric autotransporter adhesin CbpF. Journal of Oral Microbiology 11: 1565043

Brodeur BR, Boyer M, Charlebois I, Hamel JE, Couture F, Rioux MR & Martin D (2000) Identification of Group B Streptococcal Sip Protein, Which Elicits Cross-Protective Immunity. Infection & Immunity 68: 5610–5618

Carlin AF, Chang YC, Areschoug T, Lindahl G, Hurtado-Ziola N, King CC, Varki A & Nizet V (2009) Group B Streptococcus suppression of phagocyte functions by protein-mediated engagement of human Siglec-5. Journal of Experimental Medicine 206: 1691–1699

Chatellier S, Ihendyane N, Kansal RG, Khambaty F, Basma H, Norrby-Teglund A, Low DE, Mcgeer A & Kotb M (2000) Genetic Relatedness and Superantigen Expression in Group A Streptococcus Serotype M1 Isolates from Patients with Severe and Nonsevere Invasive Diseases. Infection & Immunity 68: 3523–3534

Chen SL (2019) Genomic insights into the distribution and evolution of group B streptococcus. Frontiers in Microbiology 10: DOI: 10.3389/fmicb.2019.01447

Chen T & Gotschlich EC (1996) CGM1a antigen of neutrophils, a receptor of gonococcal opacity proteins. Proc Natl Acad Sci U S A 93: 14851–14856

Conners R, Hill DJ, Borodina E, Agnew C, Daniell SJ, Burton NM, Sessions RB, Clarke AR, Catto LE, Lammie D, Wess T, Brady RL & Virji M (2008) The Moraxella adhesin UspA1 binds to its human CEACAM1 receptor by a deformable trimeric coiled-coil. EMBO Journal 27: 1779–1789

Daniel S, Nag El G, Son JPJ, Lobo FM, Htrn M, Jantscheff P, Kurok M, von Klefx S & Grunert F (1993) Determination of the Specificities of Monoclonal Antibodies Recognizing Members of the CEA Family Using a Panel of Transfectants. International Journal of Cancer 55: 303–310

Deng L, Spencer BL, Holmes JA, Mu R, Rego S, Weston TA, Hu Y, Sanches GF, Yoon S, Park N, Nagao PE, Jenkinson HF, Thornton JA, Seo KS, Nobbs AH & Doran KS (2019) The group B streptococcal surface antigen I/II protein, BspC, interacts with host vimentin to promote adherence to brain endothelium and inflammation during the pathogenesis of meningitis. PLoS Pathogens 15:

Fong JJ, Tsai CM, Saha S, Nizet V, Varki A & Buif JD (2018) Siglec-7 engagement by GBS β-protein suppresses pyroptotic cell death of natural killer cells. Proceedings of the National Academy of Sciences of the United States of America 115: 10410–10415

Gori A, Harrison OB, Mlia E, Nishihara Y, Chan JM, Msefula J, Mallewa M, Dube Q, Swarthout TD, Nobbs AH, Maiden MCJ, French N & Heyderman RS (2020) Pan-GWAS of streptococcus agalactiae highlights lineage-specific genes associated with virulence and niche adaptation. mBio 11: DOI 10.1128/mBio.00728-20

Gray-Owen SD & Blumberg RS (2006) CEACAM1: Contact-dependent control of immunity. Nature Reviews Immunology 6: 433–446

Gutbier B, Fischer K, Doehn J-M, von Lachner C, Herr C, Klaile E, Frischmann U, Singer BB, Riesbeck K, Zimmermann W, Suttorp N, Bachmann S, Bals R, Witzenrath M, Slevogt H & Lachner von C (2015) Moraxella catarrhalis induces an immune response in the murine lung that is independent of human CEACAM5 expression and long-term smoke exposure. American Journal of Physiology -Gastrointestinal and Liver Physiology 309: 250–261

Halaby DM & Mornon JPE (1998) The Immunoglobulin Superfamily: An Insight on Its Tissular, Species, and Functional Diversity. Journal of Molecular Evolution 46:389–400

Hall J, Adams NH, Bartlett L, Seale AC, Lamagni T, Bianchi-Jassir F, Lawn JE, Baker CJ, Cutland C, Heath PT, Ip M, le Doare K, Madhi SA, Rubens CE, Saha SK, Schrag S, Sobanjo-Ter Meulen A, Vekemans J & Gravett MG (2017) Maternal Disease with Group B Streptococcus and Serotype Distribution Worldwide: Systematic Review and Meta-analyses. Clinical Infectious Diseases 65: S112–S124

Heath PT (2016) Status of vaccine research and development of vaccines for GBS. Vaccine 34: 2876–2879

Hedén L-O, Frithz E & Lindahl G (1991) Molecular characterization of an IgA receptor from group B streptococci: sequence of the gene, identification of a proline-rich region with unique structure and isolation of N-terminal fragments with IgA-binding capacity. European Journal of Immunology 21: 1481–1490

Helfrich I & Singer BB (2019) Size matters: The functional role of the CEACAM1 isoform signature and its impact for NK cell-mediated killing in melanoma. Cancers 11: E356

Hill DJ, Toleman MA, Evans DJ, Villullas S, van Alphen L & Virji M (2001) The variable P5 proteins of typeable and non-typeable Haemophilus influenzae target human CEACAM1. Molecular Microbiology 39: 850–862

Hollandsworth HM, Amirfakhri S, Filemoni F, Schmitt V, Wennemuth G, Schmidt A, Hoffma RM, Singer BB & Bouvet M (2020) Anti-carcinoembryonic antigen-related cell adhesion molecule antibody for fluorescent visualization of primary colon cancer and metastases in patient-derived orthotopic xenograft mouse models. Oncotarget 11: 429–439

Hull JR, Tamura GS & Castner DG (2008) Interactions of the streptococcal C5a peptidase with human fibronectin. Acta Biomaterilia 4: 504–513.

Jauneikaite E, Kapatai G, Davies F, Gozar I, Coelho J, Bamford KB, Simone B, Begum L, Katiyo S, Patel B, Hoffman P, Lamagni T, Brannigan ET, Holmes AH, Kadhani T, Galletly T, Martin K, Lyall H, Chow Y, Godambe S, et al (2018) Serial Clustering of Late-Onset Group B Streptococcal Infections in the Neonatal Unit: A Genomic Re-evaluation of Causality. Clinical Infectious Diseases 67: 854–860

Javaheri A, Kruse T, Moonens K, Mejías-Luque R, Debraekeleer A, Asche CI, Tegtmeyer N, Kalali B, Bach NC, Sieber SA, Hill DJ, Königer V, Hauck CR, Moskalenko R, Haas R, Busch DH, Klaile E, Slevogt H, Schmidt A, Backert S, et al (2016) Helicobacter pylori adhesin HopQ engages in a virulence-enhancing interaction with human CEACAMs. Nature Microbiology 2: 16189

Khairnar V, Duhan V, Maney SK, Honke N, Shaabani N, Pandyra AA, Seifert M, Pozdeev V, Xu HC, Sharma P, Baldin F, Marquardsen F, Merches K, Lang E, Kirschning C, Westendorf AM, Häussinger D, Lang F, Dittmer U, Küppers R, et al (2015) CEACAM1 induces B-cell survival and is essential for protective antiviral antibody production. Nature Communications 6: 6217

Khairnar V, Duhan V, Patil AM, Zhou F, Bhat H, Thoens C, Sharma P, Adomati T, Friendrich SK, Bezgovsek J, Dreesen JD, Wennemuth G, Westendorf AM, Zelinskyy G, Dittmer U, Hardt C, Timm J, Göthert JR, Lang PA, Singer BB, et al (2018) CEACAM1 promotes CD8+ T cell responses and improves control of a chronic viral infection. Nature Communications 9: 2561

Königer V, Holsten L, Harrison U, Busch B, Loell E, Zhao Q, Bonsor DA, Roth A, Kengmo-Tchoupa A, Smith SI, Mueller S, Sundberg EJ, Zimmermann W, Fischer W, Hauck CR & Haas R (2016) Helicobacter pylori exploits human CEACAMs via HopQ for adherence and translocation of CagA. Nature Microbiology 2: 16188

Korotkova N, Yang Y, le Trong I, Cota E, Demeler B, Marchant J, Thomas WE, Stenkamp RE, Moseley SL & Matthews S (2008) Binding of Dr adhesins of Escherichia coli to carcinoembryonic antigen triggers receptor dissociation. Molecular Microbiology 67: 420–434

Kronvall G, Simmons A, Myhre EB & Jonsson S (1979) Specific Absorption of Human Serum Albumin, Immunoglobulin A, and Immunoglobulin G with Selected Strains of Group A-and G Streptococci. Infection and Immunity 25: 1–10

Kuespert K, Pils S & Hauck CR (2006) CEACAMs: their role in physiology and pathophysiology. Current Opinion in Cell Biology 18: 565–571

Kumar S, Stecher G & Tamura K (2016) MEGA7: Molecular Evolutionary Genetics Analysis Version 7.0 for Bigger Datasets. Molecular biology and evolution 33: 1870–1874

Kuroda M, Ohta T, Uchiyama I, Baba T, Yuzawa H, Kobayashi I, Kobayashi N, Cui L, Oguchi A, Aoki KI, Nagai Y, Lian JQ, Ito T, Kanamori M, Matsumaru H, Maruyama A, Murakami H, Hosoyama A, Mizutani-Ui Y, Takahashi NK, et al (2001) Whole genome sequencing of meticillin-resistant Staphylococcus aureus. Lancet 357: 1225–40

Larsson C, Stålhammar-Carlemalm M & Lindahl G (1996) Experimental Vaccination against Group B Streptococcus, an Encapsulated Bacterium, with Highly Purified Preparations of Cell Surface Proteins Rib and a. Infection & Immunity 64: 3518–3523

Letunic I & Bork P (2018) 20 years of the SMART protein domain annotation resource. Nucleic Acids Research 46: D493–D496

Letunic I & Bork P (2019) Interactive Tree Of Life (iTOL) v4: recent updates and new developments. Nucleic Acids Research 47: W256–W259

Lindahl G, Akerström B, Vaerman J-P & Stenberg L (1990) Characterization of an IgA receptor from group B streptococci: specificity for serum IgA. European Journal of Immunology 20: 2241–2247

Lindahl G, Stålhammar-Carlemalm M & Areschoug T (2005) Surface proteins of Streptococcus agalactiae and related proteins in other bacterial pathogens. Clinical Microbiology Reviews 18: 102–127

Michel JL, Madoff LC, Olson K, Kling DE, Kasper DL & Ausubel FM (1992) Large, identical, tandem repeating units in the C protein alpha antigen gene, bca, of group B streptococci. Proc Natl Acad Sci U S A. 89: 10060–10064

Moonens K, Hamway Y, Neddermann M, Reschke M, Tegtmeyer N, Kruse T, Kammerer R, Mejías-Luque R, Singer BB, Backert S, Gerhard M & Remaut H (2018) Helicobacter pylori adhesin HopQ disrupts trans dimerization in human CEACAMs. The EMBO Journal 37: e98665

Nagano N, Nagano Y, Nakano R, Okamoto R & Inoue M (2006) Genetic diversity of the C protein β-antigen gene and its upstream regions within clonally related groups of type la and lb group B streptococci. Microbiology 152: 771–778

Nordström T, Blom AM, Tan TT, Forsgren A & Riesbeck K (2005) Ionic Binding of C3 to the Human Pathogen Moraxella catarrhalis Is a Unique Mechanism for Combating Innate Immunity. The Journal of Immunology 175: 3628–3636

Nordström T, Movert E, Olin AI, Ali SR, Nizet V, Varki A & Areschoug T (2011) Human Siglec-5 inhibitory receptor and immunoglobulin A (IgA) have separate binding sites in streptococcal β protein. Journal of Biological Chemistry 286: 33981–33991

Patras KA & Nizet V (2018) Group B Streptococcal maternal colonization and neonatal disease: Molecular mechanisms and preventative approaches. Frontiers in Pediatrics 6: 27.

Pierce BG, Wiehe K, Hwang H, Kim BH, Vreven T & Weng Z (2014) ZDOCK server: Interactive docking prediction of protein-protein complexes and symmetric multimers. Bioinformatics 30: 1771–1773

Pietrocola G, Arciola CR, Rindi S, Montanaro L & Speziale P (2018) Streptococcus agalactiae non-pilus, cell wall-anchored proteins: Involvement in colonization and pathogenesis and potential as vaccine candidates. Frontiers in Immunology 9: 602.

Popp A, Dehio C, Grunert F, Meyer TF & Gray-Owen SD (1999) Molecular Analysis of Neisserial Opa Protein Interactions with the CEA Family of Receptors: Identification of Determinants Contributing to the Differential Specificities of Binding. Cellular Microbiology 1: 169–181

Russell-Jones GJ, Gotschlich EC & Blake MS (1984) A surface receptor specific for human IgA on group B streptococci possessing the Ibc protein antigen. Journal of Experimental Medicine 160: 1467–75.

Seale AC, Bianchi-Jassir F, Russell NJ, Kohli-Lynch M, Tann CJ, Hall J, Madrid L, Blencowe H, Cousens S, Baker CJ, Bartlett L, Cutland C, Gravett MG, Heath PT, Ip M, le Doare K, Madhi SA, Rubens CE, Saha SK, Schrag SJ, et al (2017) Estimates of the Burden of Group B Streptococcal Disease Worldwide for Pregnant Women, Stillbirths, and Children. Clinical Infectious Diseases 65: S200–S219

Shabayek S & Spellerberg B (2018) Group B streptococcal colonization, molecular characteristics, and epidemiology. Frontiers in Microbiology 9: https://doi.org/10.3389/fmicb.2018.00437

Spellerberg B, Rozdzinski E, Martin S, Weber-Heynemann J, Schnitzler N, Lütticken R, Lütticken L & Podbielski A (1999) Lmb, a Protein with Similarities to the LraI Adhesin Family, Mediates Attachment of Streptococcus agalactiae to Human Laminin. Infection & Immunity 67: 871–878.

Stålhammar-Carlemalm M, Areschoug T, Larsson C & Lindahl G (1999) The R28 protein of Streptococcus pyogenes is related to several group B streptococcal surface proteins, confers protective immunity and promotes binding to human epithelial cells. Molecular Microbiology 33: 208–219

Stålhammar-Carlemalm M, Stenberg L & Lindahl G (1993) Protein Rib: A Novel Group B Streptococcal Cell Surface Protein that Confers Protective Immunity and Is Expressed by Most Strains Causing Invasive Infections. Journal of Experimental Medicine 177: 1593–60

Tchoupa AK, Lichtenegger S, Reidl J & Hauck CR (2015) Outer membrane protein P1 is the CEACAM-binding adhesin of Haemophilus influenzae. Molecular Microbiology 98: 440–455

Tchoupa AK, Schuhmacher T & Hauck CR (2014) Signaling by epithelial members of the CEACAM family -Mucosal docking sites for pathogenic bacteria. Cell Communication and Signaling 12: 27

Tettelin H, Masignani V, Cieslewicz MJ, Donati C, Medini D, Ward NL, Angiuoli S v, Crabtree J, Jones AL, Scott Durkin A, DeBoy RT, Davidsen TM, Mora M, Scarselli M, Margarit Ros I, Peterson JD, Hauser CR, Sundaram JP, Nelson WC, Madupu R, et al (2005) Genome analysis of multiple pathogenic isolates of Streptococcus agalactiae: Implications for the microbial “‘pan-genome.’” Proc Natl Acad Sci U S A 102: 13950–5

Tong Tan T, se Nordström T, Forsgren A & Riesbeck K (2005) The Respiratory Pathogen Moraxella catarrhalis Adheres to Epithelial Cells by Interacting with Fibronectin through Ubiquitous Surface Proteins A1 and A2. Journal of Infectious Disease 192: 1029–138

Villullas S, Hill DJ, Sessions RB, Rea J & Virji M (2007) Mutational analysis of human CEACAM1: The potential of receptor polymorphism in increasing host susceptibility to bacterial infection. Cellular Microbiology 9: 329–346

Virji M, Evans D, Had^®^eld A, Grunert F, Teixeira AM & Watt SM (1999) Critical determinants of host receptor targeting by Neisseria meningitidis and Neisseria gonorrhoeae : identification of Opa adhesiotopes on the N-domain of CD66 molecules. Molecular Microbiology 34: 538–551

Virji M, Makepeace K, Ferguson DJP & Watt SM (1996) Carcinoembryonic antigens (CD66) on epithelial cells and neutrophils are receptors for Opa proteins of pathogenic neisseriae. Molecular Microbiology 22: 941–50

Wang JH (2013) The sequence signature of an Ig-fold. Protein and Cell 4: 569–572

Wang J-H & Springer TA (1998) Structural specializations of immunoglobulin superfamily members for adhesion to integrins and viruses. Immunological Reviews 163: 197–215

Watt SM, Teixeira AM, Zhou G-Q, Doyonnas R, Zhang Y, Grunert F, Blumberg RS, Kuroki M, Skubitz KM & Bates PA (2001) Homophilic adhesion of human CEACAM1 involves N-terminal domain interactions: structural analysis of the binding site. Blood 98: 1469–79

Webb B & Sali A (2016) Comparative protein structure modeling using MODELLER. Current Protocols in Bioinformatics 2016: Unit-5.6

Weng G, Wang E, Wang Z, Liu H, Zhu F, Li D & Hou T (2019) HawkDock: a web server to predict and analyze the protein-protein complex based on computational docking and MM/GBSA. Nucleic Acids Research 47: W322–W330

Winn MD, Ballard CC, Cowtan KD, Dodson EJ, Emsley P, Evans PR, Keegan RM, Krissinel EB, Leslie AGW, McCoy A, McNicholas SJ, Murshudov GN, Pannu NS, Potterton EA, Powell HR, Read RJ, Vagin A & Wilson KS (2011) Overview of the CCP4 suite and current developments. Acta Crystallographica Section D: Biological Crystallography 67: 235–242

Winstel V, Kühner P, Salomon F, Larsen J, Skov R, Hoffmann W, Peschel A & Weidenmaier C (2015) Wall teichoic acid glycosylation governs Staphylococcus aureus nasal colonization. mBio 6: e00632–15

ben Zakour NL, Venturini C, Beatson SA & Walker MJ (2012) Analysis of a Streptococcus pyogenes puerperal sepsis cluster by use of whole-genome sequencing. Journal of Clinical Microbiology 50: 2224–2228

Zhao Y, van Woudenbergh E, Zhu J, Heck AJR, van Kessel KPM, de Haas CJC, Aerts PC, van Strijp JAG & McCarthy AJ (2020) The Orphan Immune Receptor LILRB3 Modulates Fc Receptor-Mediated Functions of Neutrophils. Journal of Immunology 204: 954–966

Zhou H, Fuks A, Alcaraz G, Bolling TJ & Stanners CP (1993) Homophilic Adhesion between Ig Superfamily Carcinoembryonic Antigen Molecules Involves Double Reciprocal Bonds. Journal of Cell Biology 122: 951–960

Zimmermann L, Stephens A, Nam SZ, Rau D, Kübler J, Lozajic M, Gabler F, Söding J, Lupas AN & Alva V (2018) A Completely Reimplemented MPI Bioinformatics Toolkit with a New HHpred Server at its Core. Journal of Molecular Biology 430: 2237–2243

