## Supplemental Figures for "Bacterial protein domains with a novel Ig-like fold target human CEACAM receptors"

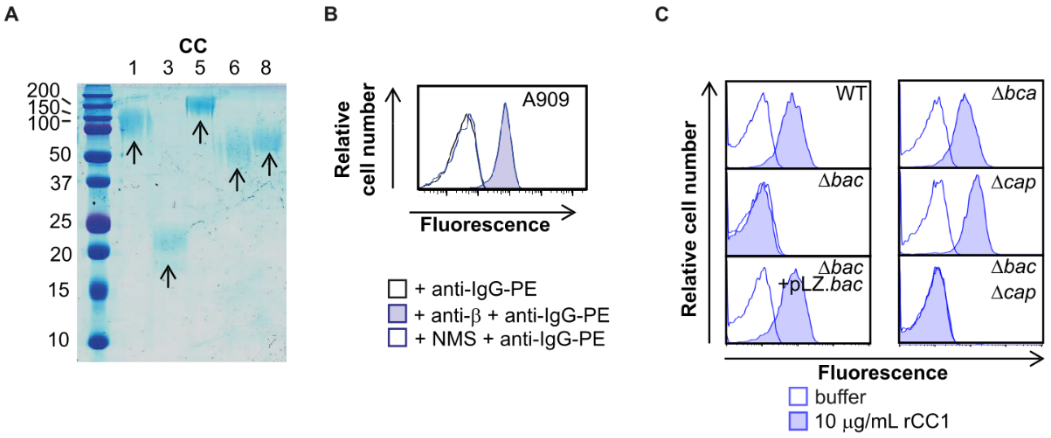

**Supplementary Figure 1: GBS binds CEACAM1 receptor through β protein. A)** Expression and purification of 6xHis tagged recombinant (r)CEACAM1 (CC1), CEACAM3 (CC3), CEACAM5 (CC5), CEACAM6 (CC6) and CEACAM8 (CC8) from a eukaryotic expression system. **B)** Mouse anti-β serum, but not normal mouse serum (NMS), detected expression of β protein in wild type A909 strain. **C)** Binding of rCC1 (10 μg/mL) to GBS strain A909 was mediated via the *bac* gene, as demonstrated by comparison of CC1 binding to wildtype, Δ*bac* and complemented strains (Δ*bac* + pLZ.*bac*). Mutation of genes encoding other surface proteins (α protein encoded by *bca*) or surface structures (capsule encoded by *cap*) did not impact rCC1 binding. Data representative of *n* = 3 replicates.

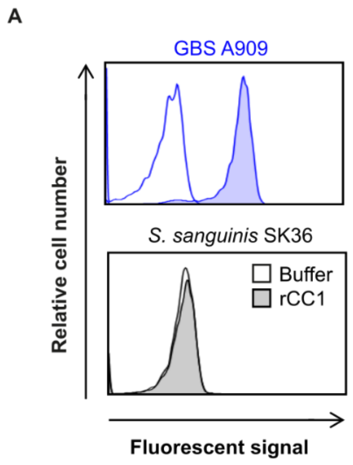

**Supplementary Figure 2: CEACAM1 does not bind to *S. sanguinis.*** Binding of rCC1 (10 µg/mL) to GBS strain A909 and *S. sanguinis* strain SK36. Data representative of *n* = 3.

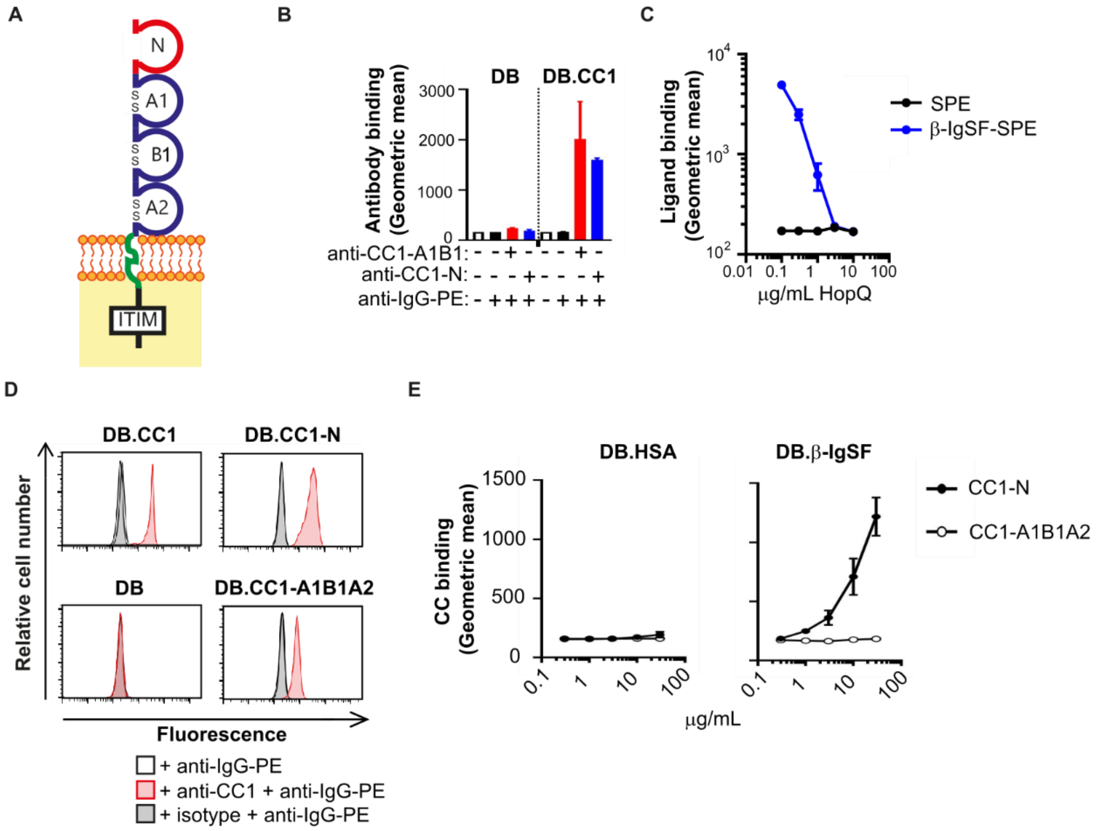

**Supplementary Figure 3: The N-terminal domain of CEACAM1 binds β-IgSF. A)** Schematic of CEACAM1 (CC1) structure. The extracellular region of the receptor is composed of the N terminal (IgV-like) domain, and the A1, B1 and A2 (IgC2) domains. The cytoplasmic tail of full length CEACAM1 contains an immunoreceptor tyrosine-based inhibitory motif (ITIM) for signalling. **B)** Control showing binding of anti-CC1-N or anti-CC1-A1B1 mAb (5 μg/mL) to dynabeads (DB) coated with CC1 or buffer. Mean and SD values are reported for *n* = 3. **C)** rHopQ inhibits binding of β-IgSF tetramers (3 μg/mL) to DB.CC1. **D)** Control showing binding of anti-CC1 mAb to DB coated with rCC1, rCC1-N and rCC1-A1B1A2 but not to controls. Data representative of *n* = 3. **E)** rCC1-N but not rCC1-A1B1A2 binds to β-IgSF-, but not HSA-, coated DB in a concentration-dependent manner. Mean and SD values are reported for *n* = 3.

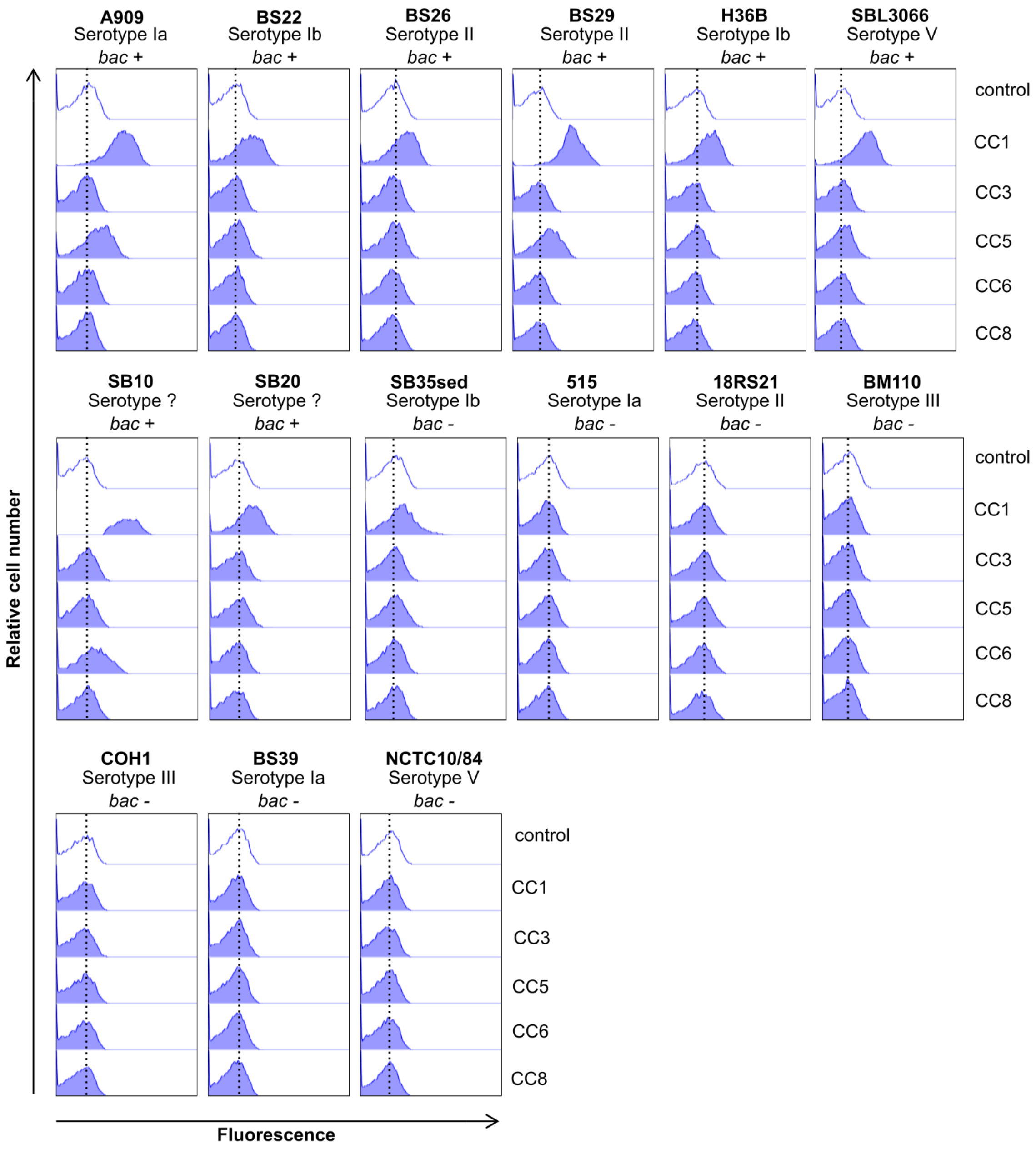

**Supplementary Figure 4: Binding of CEACAMs to GBS.** Binding rCEACAM1 (CC1), rCEACAM3 (CC3), rCEACAM5 (CC5), rCEACAM6 (CC6) and rCEACAM8 (CC8) (10 µg/mL) to a panel of GBS strains. Serotype and carriage of *bac* gene is indicated for each strain. Data representative of $n = 3$.

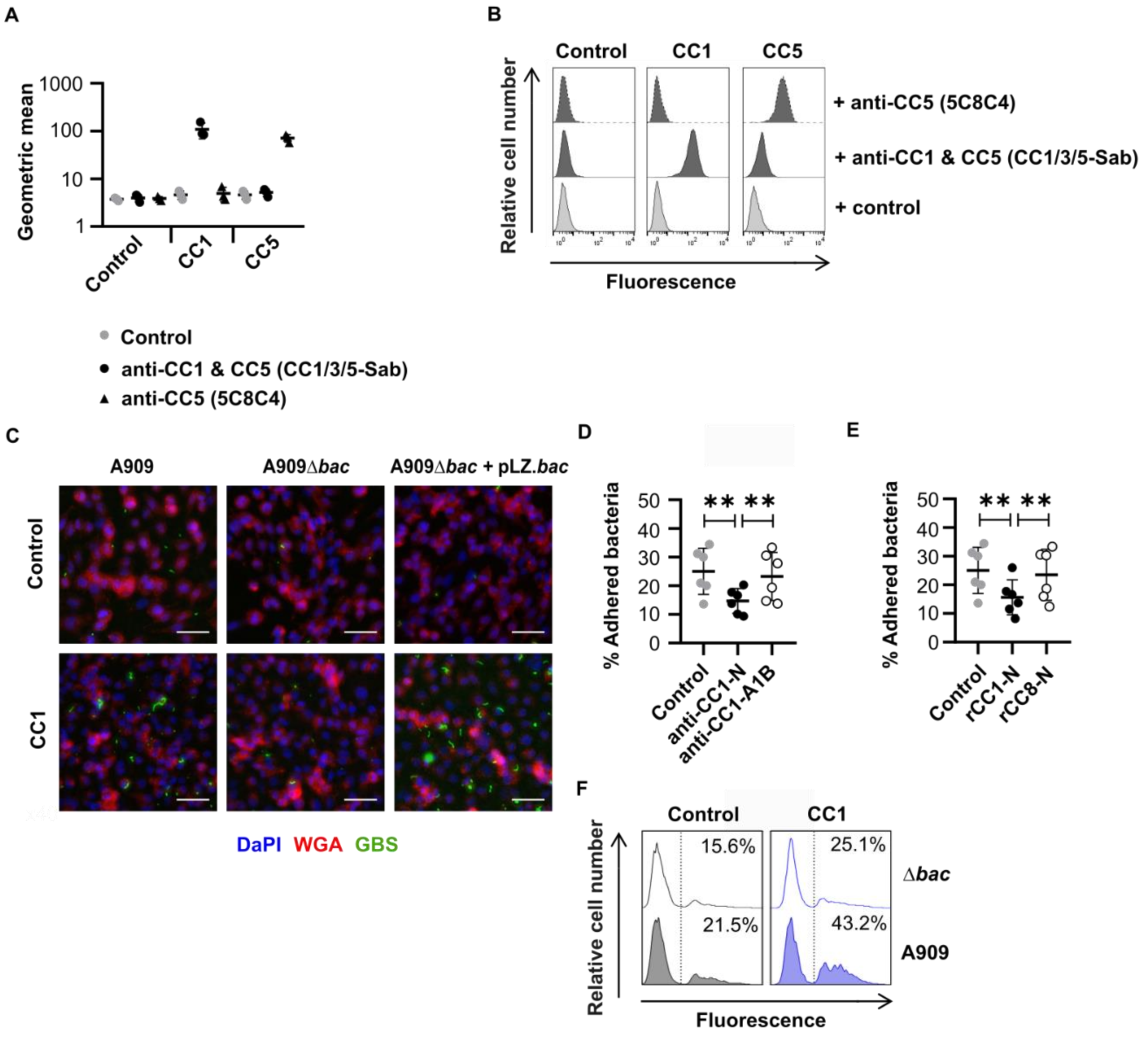

**Supplementary Figure 5: GBS exploit CEACAM1 for colonisation of epithelial cells. A and B)** Expression of human CEACAM1 (CC1) and CEACAM5 (CC5) on detached CHO cell lines, as measured by flow cytometry. CC1/3/5-Sab mAb recognises the N terminal domain of CC1, CC3 and CC5, whilst 5C8C4 mAb recognises alternative domains (A1B1A2B2A3B3) on CC5. Compiled data from *n* = 3 replicates is shown in A, and representative flow cytometry plots are show in B. **C)** Widefield microscopy imaging of FITC-labelled GBS A909, Δ*bac* or Δ*bac*+pLZ.*bac* strain adhesion to human CC1-expressing or empty vector control CHO transfectants. Cell membranes were stained with AF-647 conjugated wheat germ agglutinin (WGA)(red) and nuclei with DAPI (blue). Scale bars represent 40 μm. **D)** Adhesion of GBS A909 to CC1 expressing CHO cells was inhibited by pre-incubating cells with anti-CC1-N (clone CC1/3/5-Sab) mAb but not anti-CC1-A1B (clone B3-17) mAb. Data was analyzed by repeated One-way ANOVA with Sidak's multiple comparisons. **P*<0.05, ***P*<0.01. Mean and SD values are reported. **E)** Adhesion of GBS A909 to CC1 expressing CHO cells was inhibited by pre-incubating bacteria with 30 μg/mL rCC1-N but not rCEACAM8 (CC8)-N. Mean and SD values are reported. Data was analyzed by repeated One-way ANOVA with Sidak's multiple comparisons. **P*<0.05, ***P*<0.01. Mean and SD values are reported. **F)** Adhesion of FITC-labelled A909 strains to human CC1- expressing or control CHO transfectants at an MOI of 10. Cell lines were detached and then incubated with FITC-labelled A909 strains for 30 mins at 4ºC. Fluorescence of cells was measured by flow cytometry analysis.

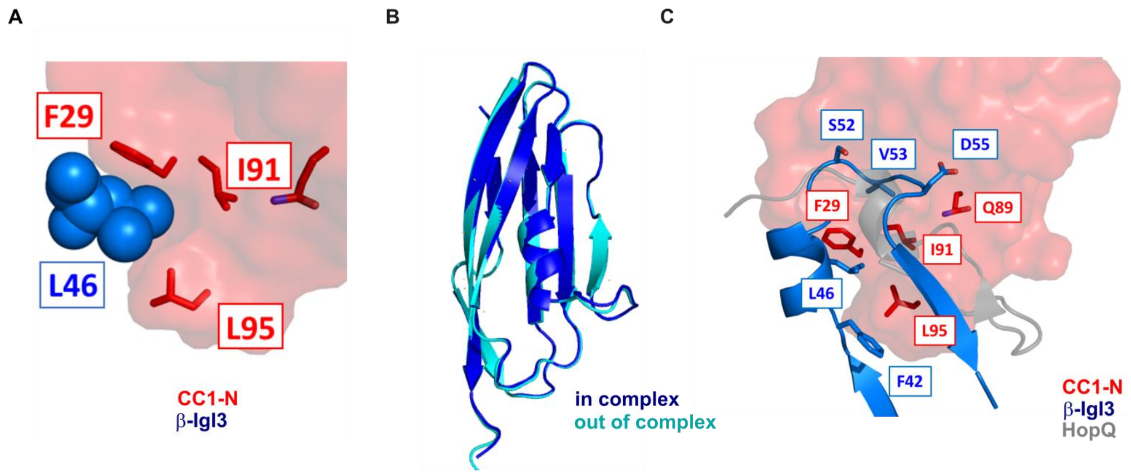

**Supplementary Figure 6: Crystal stucture of (β-IgI3)-(CEACAM1) interface. A)** Focus of the critical β-IgI3 residue L46 and interacting residues in CEACAM1 (CC1)-N. **B)** Superposition of the β-IgI3 structure out of complex (cyan) and β-IgI3 structure in complex with CC1-N complex (blue). **C)** Superposition of the β-IgI3 & CEACAM1 (CC1)-N complex (blue and red, respectively) and HopQ & CC1-N complex (grey and red, respectively).

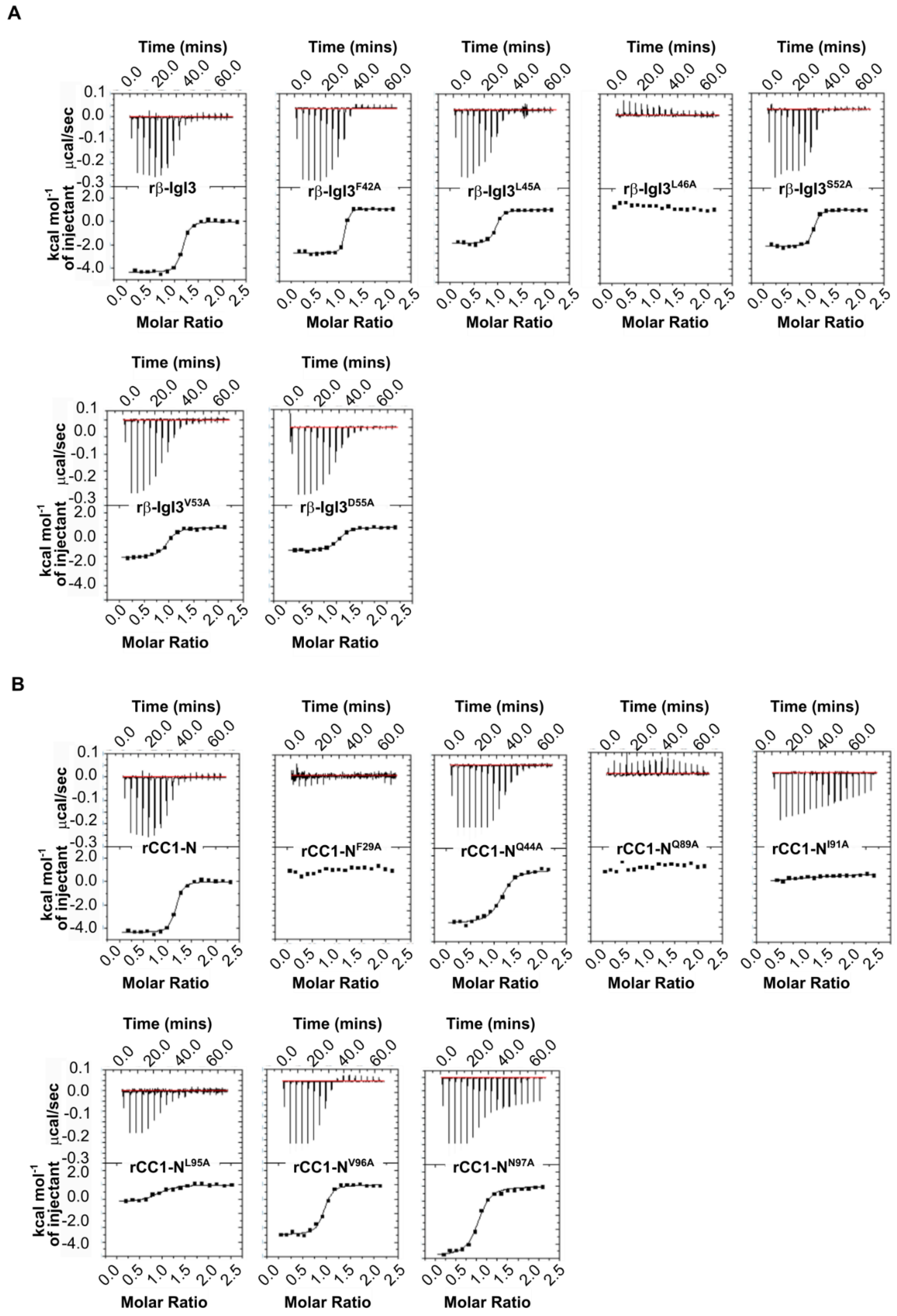

**Supplementary Figure 7: Isothermal Titration Calorimetry (ITC) binding curves. A)** rCEACM1 (CC1)-N domain and β-IgI3 mutants, and **B)** rCC1-N domain mutants and β-IgI3. Experiments were performed using an iTC200 instrument (GE Healthcare), at 25 °C with 16 injections of 2.42 µL aliquots. All data were analyzed using Origin 7.0 software.

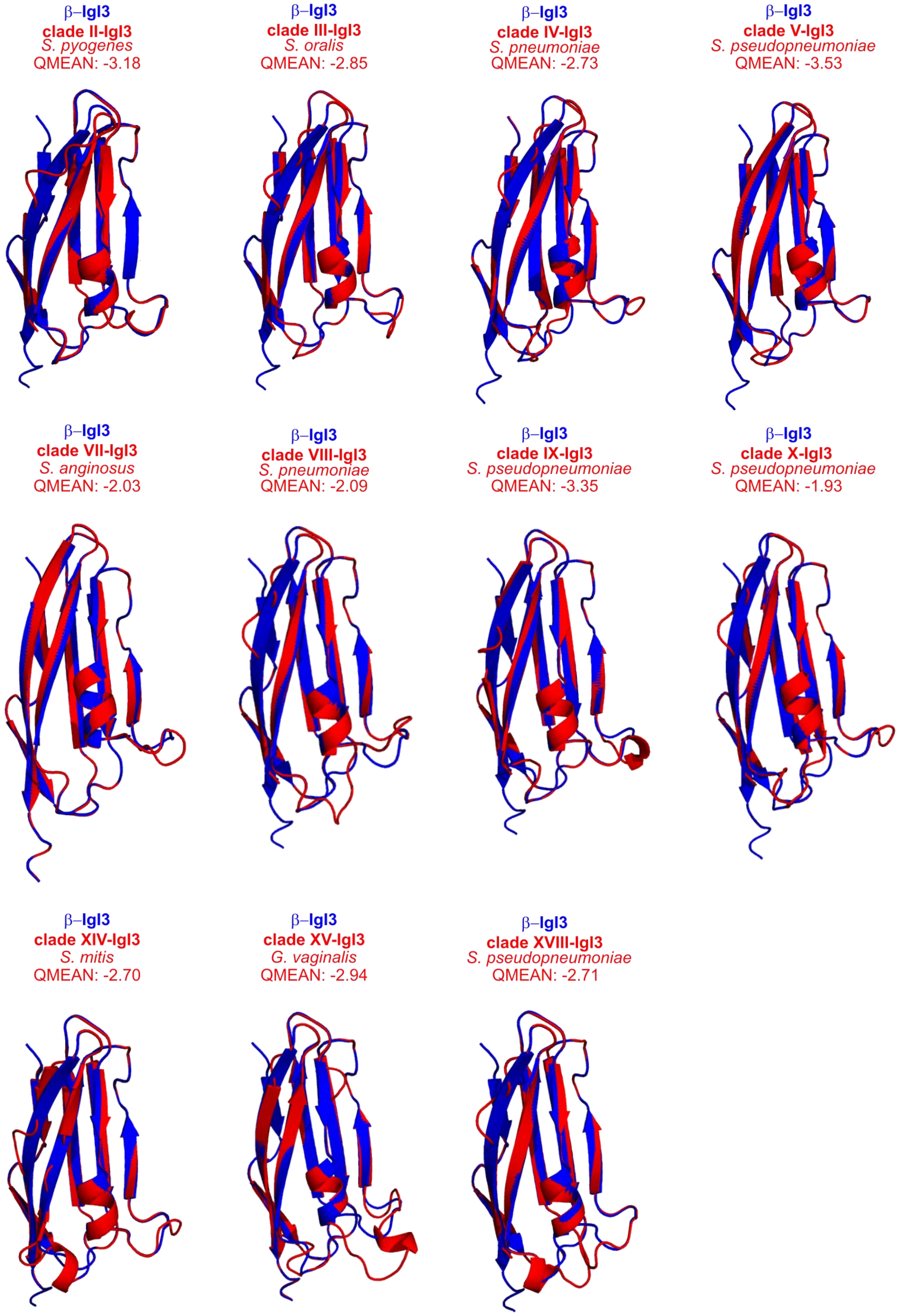

**Supplementary Figure 8: Predicted structures of IgI3 homologs.** Structure of β-IgI3 homologs was predicted using SWISS-MODEL in which the β-IgI3 was used as a template. The predicted structures of 11 sequences (red) are superimposed onto β-IgI3 structure (blue), with the clade, bacterial species and QMEAN score of the model shown.

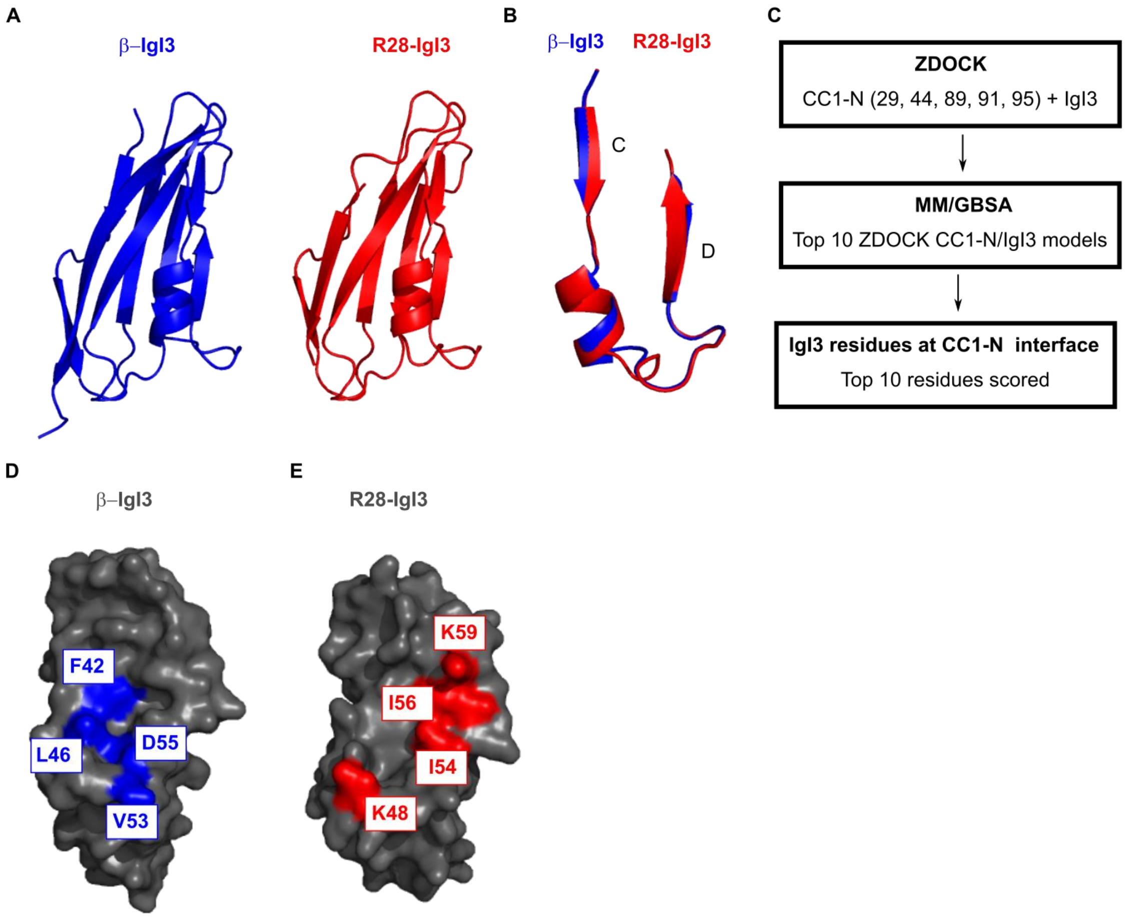

**Supplementary Figure 9: IgI3 domains bind CEACAM1 through alternative binding pockets.** **A)** Structure of β-IgI3 and predicted structure of R28-IgI3. **B)** Superposition of the known IgI3 domain from β protein (blue) onto predicted structure of IgI3 domain from R28 (red), only the *C* to *D* strand region is shown. **C)** Pipeline showing prediction of key and unfavourable IgI3 residues at the CEACAM1 binding interface. β-IgI3 and R28-IgI3 docking to CEACAM1-N (CC1-N) was simulated 50 times using ZDOCK. CC1-N residues 29, 44, 89, 91 and 95 were set as important binding residues. Each simulation was analysed by MM/GBSA analysis to quantify the free energy binding for each residue independently. **D)** Surface structure of β-IgI3 shown, with key (F42, L46 and V53) or unfavourable (D55) residues required for CC1-N binding shown in blue. **E)** Surface structure of R28-IgI3 shown, with key (K48, I54, I56 and K59) residues required for CC1-N binding shown in red.

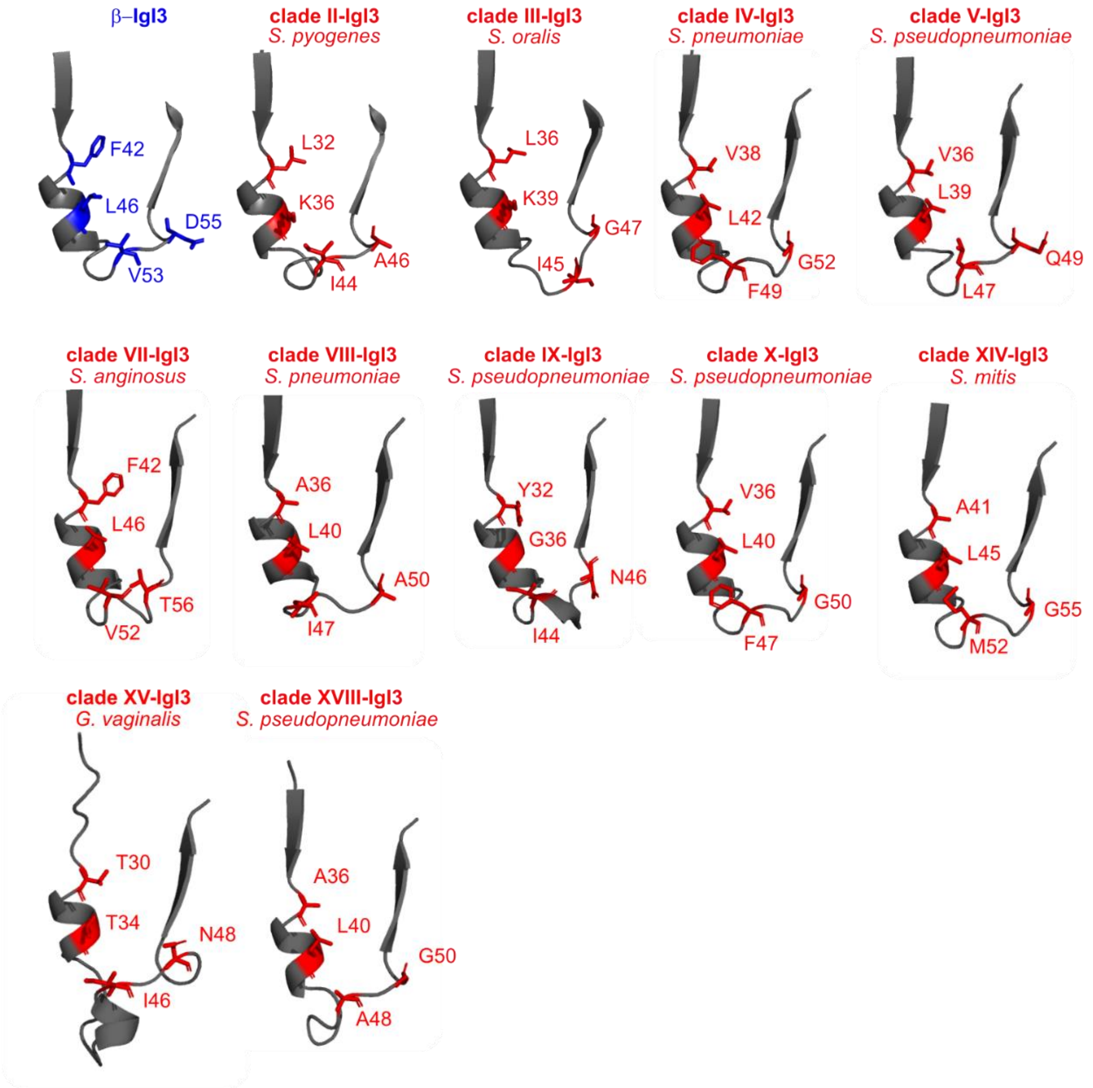

**Supplementary Figure 10: Residues in β-IgI3 required for binding CEACAM1 are not conserved in IgI3 homologs.** Close-up of the C strand, a-helix and D strand of β-IgI3 and IgI3 homologs, indicating the amino acids present at residue sites in β-IgI3 that bind CEACAM1 (CC1). The clade and bacterial species of the sequence in each model is shown.

| Species | Strain | Notes | Growth* | Reference$ |
|---|---|---|---|---|
| *Streptococcus pyogenes* (GAS) | M1 5448 | | TH-Y, 37ºC | (Chatellier *et al*, 2000) |
| | M2 | | TH-Y, 37ºC | N. van Sorge, this study |
| | M3 | | TH-Y, 37ºC | N. van Sorge, this study |
| | M6 | | TH-Y, 37ºC | N. van Sorge, this study |
| | M11 | | TH-Y, 37ºC | N. van Sorge, this study |
| | M12 | | TH-Y, 37ºC | N. van Sorge, this study |
| | M89 | | TH-Y, 37ºC | N. van Sorge, this study |
| | AL368 | | TH-Y, 37ºC | (Stålhammar-Carlemalm *et al*, 1999) |
| *Streptococcus agalactiae* (GBS) | A909 | Serotype Ia; *bac* positive | TH, 37ºC | (Michel *et al*, 1992) |
| | A909Δ*bac* | | TH, 37ºC | (Areschoug *et al*, 2002b) |
| | A909Δ*bac + pLZ.bac* | | TH + k + s, 37ºC | (Areschoug *et al*, 2002b) |
| | A909Δ*bca* | | TH + k, 37ºC | E. Movert, T. Areschoug, Lund University |
| | A909Δ*cap* | | TH + k, 37ºC | E. Movert, T. Areschoug, Lund University |
| | A909Δ*bac*Δ*cap* | | TH + k, 37ºC | E. Movert, T. Areschoug, Lund University |
| | BS39 | Serotype Ia; *bac* negative; invasive isolate | TH, 37ºC | G. Lindahl, this study |
| | 515 | Serotype Ia; *bac* negative | TH, 37ºC | (Areschoug *et al*, 2002a) |
| | BS22 | Serotype Ib; *bac* positive | TH, 37ºC | (Areschoug *et al*, 2002a) |
| | H36B | Serotype Ib; *bac* positive | TH, 37ºC | ATCC #BAA-1174 |
| | BS29 | Serotype II; *bac* positive; invasive isolate | TH, 37ºC | G. Lindahl, this study |
| | 18RS21 | Serotype II; *bac* negative | TH, 37ºC | ATCC #BAA1175 |
| | BM110 | Serotype III; *bac* negative | TH, 37ºC | (Stålhammar-Carlemalm *et al*, 1993) |
| | COH1 | Serotype III; *bac* negative | TH, 37ºC | ATCC #BAA-1176 |
| | SBL3066 | Serotype V; *bac* positive; invasive isolate | TH, 37ºC | G. Lindahl, this study |
| | NCTC10/84 | Serotype V; *bac* negative | TH, 37ºC | ATCC #49447 |
| | SB35 | *bac* positive | TH, 37ºC | (Stålhammar-Carlemalm *et al*, 1993) |
| | SB10 | *bac* positive | TH, 37ºC | (Areschoug *et al*, 2002a) |
| | SB20 | Serotype III; *bac* positive | TH, 37ºC | (Areschoug *et al*, 2002a) |
| | BS26 | *bac* positive | TH, 37ºC | (Areschoug *et al*, 2002a) |
| | PHEGBS0159 | *alp3* positive | TH, 37ºC | (Jauneikaite *et al*, 2018) |
| *Streptococcus zooepidemicus* | CS2 | Invasive infection | TH, 37ºC | G. Lindahl, this study |
| | CS3 | Invasive infection | TH, 37ºC | G. Lindahl, this study |

| | | | | |
|---|---|---|---|---|
| (GCS) | CS4 | Invasive infection | TH, 37ºC | G. Lindahl, this study |
| | CS7 | Invasive infection | TH, 37ºC | G. Lindahl, this study |
| | CS8 | Invasive infection | TH, 37ºC | G. Lindahl, this study |
| | L31 | Invasive infection | TH, 37ºC | G. Lindahl, this study |
| | L32 | Invasive infection | TH, 37ºC | G. Lindahl, this study |
| *Streptococcus dysgalactiae* (GGS) | G148 | | TH, 37ºC | (Kronvall *et al*, 1979) |
| | GS1 | Invasive infection | TH, 37ºC | G. Lindahl, this study |
| | GS2 | Invasive infection | TH, 37ºC | G. Lindahl, this study |
| | GS3 | Invasive infection | TH, 37ºC | G. Lindahl, this study |
| | GS4 | Invasive infection | TH, 37ºC | G. Lindahl, this study |
| | GS5 | Invasive infection | TH, 37ºC | G. Lindahl, this study |
| | GS6 | Invasive infection | TH, 37ºC | G. Lindahl, this study |
| | GS7 | Invasive infection | TH, 37ºC | G. Lindahl, this study |
| | GS8 | Invasive infection | TH, 37ºC | G. Lindahl, this study |
| | GS9 | Invasive infection | TH, 37ºC | G. Lindahl, this study |
| | L33 | Invasive infection | TH, 37ºC | G. Lindahl, this study |
| | L34 | Invasive infection | TH, 37ºC | G. Lindahl, this study |
| *Streptococcus pneumoniae* | PBCN22 | | TH, 37ºC | UMC Utrecht, this study |
| | D39 | | TH, 37ºC | NCTC #7466 |
| | TIGR4 | | TH, 37ºC | BAA-334 |
| | PBCN57 | | TH, 37ºC | UMC Utrecht, this study |
| | PBCN79 | | TH, 37ºC | UMC Utrecht, this study |
| | PBCN24 | | TH, 37ºC | UMC Utrecht, this study |
| | PBCN133 | | TH, 37ºC | UMC Utrecht, this study |
| *Enterococcus faecium* | E8284 | | TH, 37ºC | (Arredondo-Alonso *et al*, 2020) |
| | E4413 | | TH, 37ºC | (Arredondo-Alonso *et al*, 2020) |
| | E656 | | TH, 37ºC | (Arredondo-Alonso *et al*, 2020) |
| | E4227 | | TH, 37ºC | (Arredondo-Alonso *et al*, 2020) |
| | E7313 | | TH, 37ºC | (Arredondo-Alonso *et al*, 2020) |
| | E7098 | | TH, 37ºC | (Arredondo-Alonso *et al*, 2020) |
| *Enterococcus faecalis* | E02500 | Human blood isolate | TH, 37ºC | UMC Utrecht, this study |
| | E02504 | Human blood isolate | TH, 37ºC | UMC Utrecht, this study |
| | E02608 | Human blood isolate | TH, 37ºC | UMC Utrecht, this study |

| | | | | |
|---|---|---|---|---|
| | E02835 | Human blood isolate | TH, 37$^{o}$C | UMC Utrecht, this study |
| | E4877 | Human faecal isolate | TH, 37$^{o}$C | UMC Utrecht, this study |
| | E6568 | Human faecal isolate | TH, 37$^{o}$C | UMC Utrecht, this study |
| *Staphylococcus aureus* | MW2 | | TSB, 37$^{o}$C | ATCC #BAA-1707 |
| | MRSA252 | | TSB, 37$^{o}$C | ATCC #BAA-1720 |
| | PS66 | | TSB, 37$^{o}$C | U. Bläsi, Vienna |
| | 80286 | | TSB, 37$^{o}$C | (Winstel *et al*, 2015) |
| | USA300 | | TSB, 37$^{o}$C | ATCC #BAA-1556 |
| | N315 | | TSB, 37$^{o}$C | (Kuroda *et al*, 2001) |
| | Newman | | TSB, 37$^{o}$C | (Baba *et al*, 2008) |

**Supplementary Table 1. Bacterial strains used in this study.** * Growth media as follows, Todd-Hewitt (TH) broth, Todd-Hewitt + 0.6% yeast (TH-Y) broth, Tryptic Soy (TS) broth. Media was supplemented with 500 μg/ml kanamycin (k) and/or 70 μg/ml spectinomycin (s). [$] Strain names available from American Type Culture Collection (ATCC) or National Collection of Type Cultures (NCTC).

| Vector | Protein | Expression system |
|---|---|---|
| pcDNA3.4.CEACAM1 | rCEACAM1-His | Expi293F |
| pcDNA3.4.CEACAM3 | rCEACAM3-His | Expi293F |
| pcDNA3.4.CEACAM5 | rCEACAM5-His | Expi293F |
| pcDNA3.4.CEACAM6 | rCEACAM6-His | Expi293F |
| pcDNA3.4.CEACAM8 | rCEACAM8-His | Expi293F |
| pRSET-C-CEACAM1N | rCEACAM1-N-His | *E. coli* RG |
| pRSET-C-CEACAM1NΔF29A | rCEACAM1-NΔF29A-His | *E. coli* RG |
| pRSET-C-CEACAM1NΔQ44A | rCEACAM1-NΔQ44A-His | *E. coli* RG |
| pRSET-C-CEACAM1NΔA49V | rCEACAM1-NΔA49V-His | *E. coli* RG |
| pRSET-C-CEACAM1NΔQ89A | rCEACAM1-NΔQ89A-His | *E. coli* RG |
| pRSET-C-CEACAM1NΔL95A | rCEACAM1-NΔL95A-His | *E. coli* RG |
| pRSET-C-CEACAM1NΔV96A | rCEACAM1-NΔV96A-His | *E. coli* RG |
| pRSET-C-CEACAM1NΔN97A | rCEACAM1-NΔN97A-His | *E. coli* RG |
| pRSET-C-CEACAM1A1B1A2 | rCEACAM1-A1B1A2-His | *E. coli* RG |
| pRSET-C-CEACAM3N | rCEACAM3-N-His | *E. coli* RG |
| pRSET-C-CEACAM5N | rCEACAM5-N-His | *E. coli* RG |
| pRSET-C-CEACAM6N | rCEACAM6-N-His | *E. coli* RG |
| pRSET-C-CEACAM8N | rCEACAM8-N-His | *E. coli* RG |
| pET21d-CEACAM1N | rCEACAM1-N | *E. coli* RG |
| pET21d-CEACAM1NΔF29A | rCEACAM1-NΔF29A | *E. coli* RG |
| pET21d-CEACAM1NΔQ44A | rCEACAM1-NΔQ44A | *E. coli* RG |
| pET21d-CEACAM1NΔA49V | rCEACAM1-NΔA49V | *E. coli* RG |
| pET21d-CEACAM1NΔQ89A | rCEACAM1-NΔQ89A | *E. coli* RG |
| pET21d-CEACAM1NΔL95A | rCEACAM1-NΔL95A | *E. coli* RG |
| pET21d-CEACAM1NΔV96A | rCEACAM1-NΔV96A | *E. coli* RG |
| pET21d-CEACAM1NΔN97A | rCEACAM1-NΔN97A | *E. coli* RG |
| pRSET-C-B6N | rB6N-His | *E. coli* RG |
| pRSET-C-IgABR | rIgABR-His | *E. coli* RG |
| pRSET-C-B6C | rB6C-His | *E. coli* RG |
| pRSET-C-β-IgSF / β-IgI3 | rβIgSF-His, β-IgI3-His | *E. coli* RG |
| | rβIgI3ΔF42A-His | *E. coli* RG |
| | rβIgI3ΔL45A-His | *E. coli* RG |
| | rβIgI3ΔL46A-His | *E. coli* RG |
| | rβIgI3ΔS52A-His | *E. coli* RG |
| | rβIgI3ΔV53A-His | *E. coli* RG |
| | rβIgI3ΔD55A-His | *E. coli* RG |
| pRSET-C-β75KN | rβ75KN-His | *E. coli* RG |
| pRSET-C-R28-IgI3 | rR28-IgI3-His | *E. coli* RG |

**Supplementary Table 2: Expression vectors constructs.** Open reading frames (ORFs) coding the extracellular domains of CEACAMs were cloned into pcDNA3.4 vectors, and proteins were expressed in Expi293F cells. ORFs coding the N domains of CEACAMs, the A1B1A2 domain of CEACAM1 and β protein domains were cloned into pRSET-C vectors, and proteins were expressed in *E. coli*.

| | **$K_D$ (nM)** | **ΔH (kcal $mol^{-1}$)** | **TΔS (kcal $mol^{-1}$)** |
|---|---|---|---|
| **β-IgSF** | | | |
| rCC1-N | 96±2 | -4.7±0.3 | +4.9 |
| rCC3-N | No Binding Observed | | |
| rCC5-N | 152±27 | -2.2±0.1 | +7.1 |
| rCC6-N | No Binding Observed | | |
| rCC8-N | No Binding Observed | | |

**Supplementary Table 3: Isothermal Titration Calorimetry (ITC) binding curves constants and thermodynamic paramters for CEACAM-N and β-IgSF interactions.** Experiments were performed using β-IgI3 and N domains of (r)CEACAM1 (CC1), CEACAM3 (CC3), CEACAM5 (CC5), CEACAM6 (CC6) and CEACAM8 (CC8) on an iTC200 instrument (GE Healthcare), at 25 °C with 16 injections of 2.42 µL aliquots. All data were analyzed using Origin 7.0 software.

| | **β-IgI3-CC1-N** |
|---|---|
| **Data collection** | |
| Space group | I4122 |
| Cell dimensions | |
| *a*, *b*, *c* (Å) | 131.68 131.68 256.91 |
| α, β, γ (°) | 90.00 90.00 90.00 |
| Resolution (Å) | 48.57-2.95 |
| $R_{\text{merge}}$ | 0.140 (9.455) |
| $R_{p.i.m}$ | 0.046 (3.134) |
| $I / \sigma I$ | 13.2 (0.5) |
| Completeness (%) | 99.9 (99.9) |
| Redundancy | 19.2 (19.1) |
| $CC_{1/2}$ | 0.999 (0.592) |
| **Refinement** | |
| Resolution (Å) | 48.57-2.95 |
| No. reflections | 22918 |
| $R_{\text{work}} / R_{\text{free}}$ | 23.1/26.5 |
| No. atoms | |
| Protein | 3311 |
| Water | 2 |
| Ligand | 37 |
| *B*-factors | |
| Protein | 151.5 |
| Water | 105.1 |
| Ligands | 179.0 |
| R.m.s. deviations | |
| Bond lengths (Å) | 0.004 |
| Bond angles (°) | 1.473 |

**Supplementary Table 4**. **Data collection and refinement statistics for β-IgI3 and CEACAM1 (CC1)-N**. Values in parentheses are for highest-resolution shell.

| β-IgI3 atom | Distance (Å) | CC1-N atom |
|---|---|---|
| D41 [OD1] | 3.92 | S93 [CB] |
| F42 [CB] | 3.44 | L95 [CD2] |
| F42 [CG] | 2.96 | L95 [CD2] |
| F42 [CD1] | 2.87 | L95 [CD2] |
| F42 [CE1] | 3.31 | L95 [CD2] |
| F42 [CZ] | 3.76 | L95 [CD2] |
| F42 [CD2] | 3.41 | L95 [CD2] |
| F42 [CE2] | 3.79 | L95 [CD2] |
| L46 [CB] | 3.63 | F29 [CZ] |
| L46 [CB] | 3.65 | F29 [CD2] |
| L46 [CB] | 3.44 | F29 [CE2] |
| L46 [CB] | 3.99 | F29 [CE1] |
| L46 [CG] | 3.99 | F29 [CD2] |
| L46 [CD1] | 3.64 | F29 [CB] |
| L46 [CD1] | 3.44 | F29 [CG] |
| L46 [CD1] | 3.11 | F29 [CG] |
| L46 [CD1] | 3.78 | F29 [CE2] |
| L46 [CD2] | 3.61 | L95 [CD1] |
| S52 [CB] | 3.63 | T56 [OG1] |
| S52 [CB] | 3.56 | T56 [CB] |
| S52 [CB] | 3.92 | T56 [CG2] |
| V53 [CB] | 3.94 | S32 [OG] |
| V53 [CG1] | 3.16 | I91 [CG1] |
| V53 [CG1] | 3.36 | S32 [OG] |
| V53 [CG1] | 3.66 | I91 [CD1] |
| V53 [CG1] | 3.94 | S32 [CB] |
| V53 [CG2] | 3.84 | I91 [CD1] |
| S54 [CB] | 3.87 | Y34 [OH] |
| S54 [CB] | 3.94 | Q44 [CD] |
| S54 [CB] | 3.81 | Q44 [OE1] |
| S54 [CB] | 3.74 | Q44 [NE2] |
| D55 [OD1] | 3.82 | Y34 [OH] |
| D55 [OD1] | 3.57 | Y34 [CE2] |
| T59 [CB] | 3.86 | L95 [CB] |
| T59 [OG1] | 3.39 | L95 [CB] |
| T59 [CG2] | 3.43 | L95 [CB] |
| T59 [CG2] | 3.43 | L95 [CG] |
| Y61 [OH] | 3.75 | D94 [CB] |
| Y61 [CE2] | 3.30 | V96 [CG1] |
| Y61 [CD2] | 3.93 | V96 [CG1] |

**Supplementary Table 5. Contact sites in (β-IgI3)(CEACAM1-N) complex.** β-IgI3 atoms within 4.0Å of CEACAM1-N (CC1-N) atoms are displayed as calculated using Ncont from the CCP4 suite of programs.

| | $K_D$ (nM) | ΔH (kcal mol$^{-1}$) | TΔS (kcal mol$^{-1}$) |
|---|---|---|---|
| **Binding to rCC1-N** | | | |
| β-IgI3 | 96±2 | -4.7±0.3 | +4.9 |
| β-IgI3$^{F42A}$ | 16±15 | -6.3±0.2 | +5.2 |
| β-IgI3$^{L45A}$ | 234±16 | -4.7±0.1 | +4.3 |
| β-IgI3$^{L46A}$ | No Binding Observed | | |
| β-IgI3$^{S52A}$ | 85±20 | -4.8±0.1 | +4.9 |
| β-IgI3$^{V53A}$ | 562±44 | -4.3±0.0 | +4.3 |
| β-IgI3$^{D55A}$ | 690±52 | -3.1±0.1 | +5.3 |
| **Binding to β-IgI3** | | | |
| rCC1-N | 96±2 | -4.7±0.3 | +4.9 |
| rCC1-N$^{F29A}$ | No Binding Observed | | |
| rCC1-N$^{Q44A}$ | 996±116 | -7.6±0.1 | +0.6 |
| rCC1-N$^{Q89A}$ | No Binding Observed | | |
| rCC1-N$^{I91A}$ | No Binding Observed | | |
| rCC1-N$^{L95A}$ | 1350±460 | -2.3±0.1 | +5.7 |
| rCC1-N$^{V96A}$ | 370±4 | -7.0±0.1 | +1.8 |
| rCC1-N$^{N97A}$ | 490±120 | -9.2±0.4 | -0.5 |

**Supplementary Table 6: Isothermal Titration Calorimetry (ITC) binding curves constants and thermodynamic paramters for CEACAM1 (CC1)-N and β-IgI3 mutants.** Experiments were performed using an iTC200 instrument (GE Healthcare), at 25 °C with 16 injections of 2.42 µL aliquots. All data were analyzed using Origin 7.0 software.
